# The metabolic kinase LKB1 shapes the enteric nervous system by mitigating the oxidative stress and p53 activity

**DOI:** 10.1101/2025.06.10.658770

**Authors:** Anthony Lucas, Florence Appaix, Jordan Allard, Marie Mével-Aliset, Anca G. Radu, Florence Fauvelle, Bertrand Favier, Alexei Grichine, Jochen Maurer, Pierre Hainaut, Attardi Laura, Marc Billaud, Sakina Torch, Chantal Thibert

**Author notes:** Contributed equally to the work (Anthony Lucas, Florence Appaix). corresponding and co-senior authors: to whom correspondence should be sent.

## Abstract

The enteric nervous system (ENS) comprises ganglia of neurons and glial cells derived from migratory multipotent neural crest cells. While the molecular mechanisms of ENS development are well-studied, the involvement of metabolic processes has received less attention. We previously showed that the tumor suppressor kinase LKB1 is essential for the trophic maintenance of postnatal ENS. Here we examined LKB1’s role in ENS formation using a genetically engineered mouse model that conditionally inactivates *Lkb1* in neural crest progenitors during gut invasion. We conducted a comprehensive phenotyping of the ENS through histology and 3D imaging of cleared tissue, combining lightsheet microscopy with adaptive optics confocal microscopy. We found that *Lkb1* loss impairs early neuronal differentiation, followed by glial degeneration, leading to hypoganglionosis and compromised digestive tissue integrity. Metabolite profiling of digestive tracts revealed an increase of oxidative stress upon *Lkb1* ablation. *In vitro, Lkb1* knockdown induced oxidative stress in neural crest progenitors and their glial derivatives, causing DNA damage and p53 activation. Ablation of p53 rescued glial specification under these conditions. *In vivo*, hyperphosphorylation of p53 was also observed; however, deletion of *p53* alleles in *Lkb1* mutants did not restore enteric neurons number. Instead, it improved axonal fiber organization and partially rescued digestive tissue integrity. These findings establish LKB1 as a key metabolic regulator on both the development and maintenance of the ENS, suggesting that aberrant LKB1 signaling may contribute to human enteric glioneuropathies.

**Highlights:**

- LKB1 loss in enteric progenitors results in extensive hypoganglionosis and disrupts gut tissue homeostasis.
- During embryogenesis, LKB1 shapes enteric ganglia by sequentially regulating neuronal differentiation and preserving glial cells, partly by limiting oxidative stress and p53 activity.
- These findings establish LKB1 as a critical regulator of neural crest cell formation, highlighting its multifaceted roles and potential pathological implications in digestive neuropathies.

## Introduction

The enteric nervous system (ENS), a subdivision of the peripheral nervous system, autonomously controls digestive motility and secretion, local blood flow, immune response and microbiota interaction, and also plays a central role in the gut-brain axis (Sharkey and Mawe 2023). Accordingly, ENS dysfunction is not only associated with enteric neuropathies but also contributes to neurodegenerative diseases and neurodevelopmental disorders (Niesler et al. 2021; Margolis et al. 2021; Holland et al. 2021). Given the rising occurrence of these pathologies, a better understanding of the various factors that shape ENS structure and function is essential.

The ENS consists of two concentric, interconnected layers of ganglia: the myenteric and submucosal plexuses, embedded within the smooth muscle layers (*tunica muscularis*) of the digestive tract (Sharkey and Mawe 2023). Neurons and glial cells of the ENS arise from neural crest cells (NCCs), which are multipotent migratory embryonic cells that delaminate from the dorsal part of the neural tube (Thibert et al. 2023). This transient population of cells reach numerous organs, giving rise to a diverse array of derivatives, including bones and cartilages of the face, melanocytes in the skin, neuroendocrine cells, sensory neurons and Schwann cells in the peripheral nerves, as well as enteric neurons and glial cells (Perera and Kerosuo 2021; Rao and Gershon 2018). The NCCs that form enteric ganglia originate from vagal and sacral neural crest subpopulations, supplemented by neural crest derivatives known as Schwann cell precursors, which are closely associated with the vagal nerve and sympathetic fibers that provide extrinsic innervation to the gut via the mesentery (Kameneva et al. 2021).

NCC development relies on well-defined gene regulatory networks that govern the neural crest lineage trees (Kastriti et al. 2022; Erickson et al. 2023; Zurkirchen and Sommer 2017; Lasrado et al. 2017). However, our understanding of the role of metabolic regulation in NCC development has only recently emerged, with aerobic glycolysis playing a key role in cranial NCCs, while trunk NCCs rely on glucose oxidation (Bhattacharya et al. 2020; Nekooie Marnany et al. 2023; Barriga et al. 2018). In addition, we have previously uncovered the critical and multifaceted role of the metabolic regulator LKB1 in NCC formation (Creuzet et al. 2016; Radu et al. 2019; Thibert et al. 2023). LKB1 is a master serine/threonine kinase that governs cell growth, polarity and energy metabolism by activating 14 downstream kinases, including the energy sensor AMP-activated protein kinase (AMPK) (Kullmann and Krahn 2018). To investigate the role of LKB1 in NCCs, we generated two genetically engineered mouse models that enable selective and spatiotemporal inactivation of *Lkb1* in either post-delaminating NCCs (from ∼E8) or post-migrating NCCs (from ∼E10.5). This approach allowed us to dissect temporal LKB1’s functions throughout neural crest development.

Inactivation of *Lkb1* in post-migrating NCCs led to lethality around weaning, with mutant mice displaying a complex neurocristopathy phenotype, including coat depigmentation, peripheral neuropathy, and intestinal pseudo-obstruction (Radu et al. 2019). Histological analysis revealed a progressive postnatal degeneration of the ENS, leading to hypoganglionosis. Specification of neural crest-derived glial cells was found to depend on LKB1-mediated regulation of pyruvate-alanine interconversion coupled with mTOR signaling. Treatment with 5-Aminoimidazole-4-carboxamide riboside (AICAR), an AMP precursor, partially rescued motor defects and enteric ganglia loss (Radu et al. 2019).

In contrast, early *Lkb1* inactivation in pre-migrating NCCs led to perinatal lethality (Creuzet et al. 2016), underscoring LKB1’s essential role in vertebrate head formation through the regulation of cephalic NCCs delamination, migration, survival, and differentiation (Creuzet et al. 2016). This process involved the AMPK/Rho/Rock pathway to control actin dynamics (Creuzet et al. 2016). Moreover, early Lkb1 inactivation resulted in additional defects across other neural crest derivatives, indicating a broader neurocristopathy phenotype comparable to that observed with later-stage Lkb1 loss.

In this study, we investigated the functional consequences of *Lkb1* ablation in enteric progenitors and ENS formation using our genetic model targeting *Lkb1* ablation specifically in pre-migrating NCCs. Comprehensive phenotyping was performed using classical histology and 3D imaging of cleared tissues, combining lightsheet and adaptive optics confocal microscopy. Our findings demonstrate that Lkb1 loss sequentially impairs early neuronal differentiation and later glial cell maintenance, leading to hypoganglionosis and impaired digestive tissue integrity. Metabolite profiling on digestive tracts revealed elevated oxidative stress following *Lkb1* ablation. *In vitro, Lkb1* knockdown similarly induced oxidative stress in neural crest progenitors and glial derivatives, causing DNA damage and p53 activation. Under these conditions, p53 ablation rescued glial specification. *In vivo*, we also observed hyperphosphorylation in Lkb1 mutants; however, deletion of *p53* alleles did not restore enteric neurons numbers. Instead, it improved axonal fiber organization and partially rescued digestive tissue architecture.

Together, these findings establish LKB1 as a central regulator of ENS development and maintenance, acting in part by limiting oxidative stress and preventing p53 hyperactivation. They further suggest that dysregulation of the LKB1 pathway may play a pathogenic role in enteric glioneuropathies.

## Results

### Lkb1 signaling in vagal neural crest cells is essential for enteric ganglia formation

To investigate the role of LKB1 in enteric nervous system development, we conducted an extensive phenotyping of a genetically engineered mouse model that enables the conditional deletion of *Lkb1* floxed alleles in pre-migratory NCCs from embryonic day E8.5 (Fig1A). This ablation is initiated by Cre recombinase expression driven by the *HtPA* (human tissue plasminogen activator) promotor, with Cre expression traceable via the *LacZ* reporter (R26R). This model, referred to as LRH mice, leads to perinatal lethality and marked craniofacial malformations (Creuzet et al. 2016). It also affects overall digestive tract homeostasis (Figure 1B), suggestive of defective vagal neural crest progenitors and derivatives.

**Figure 1.**
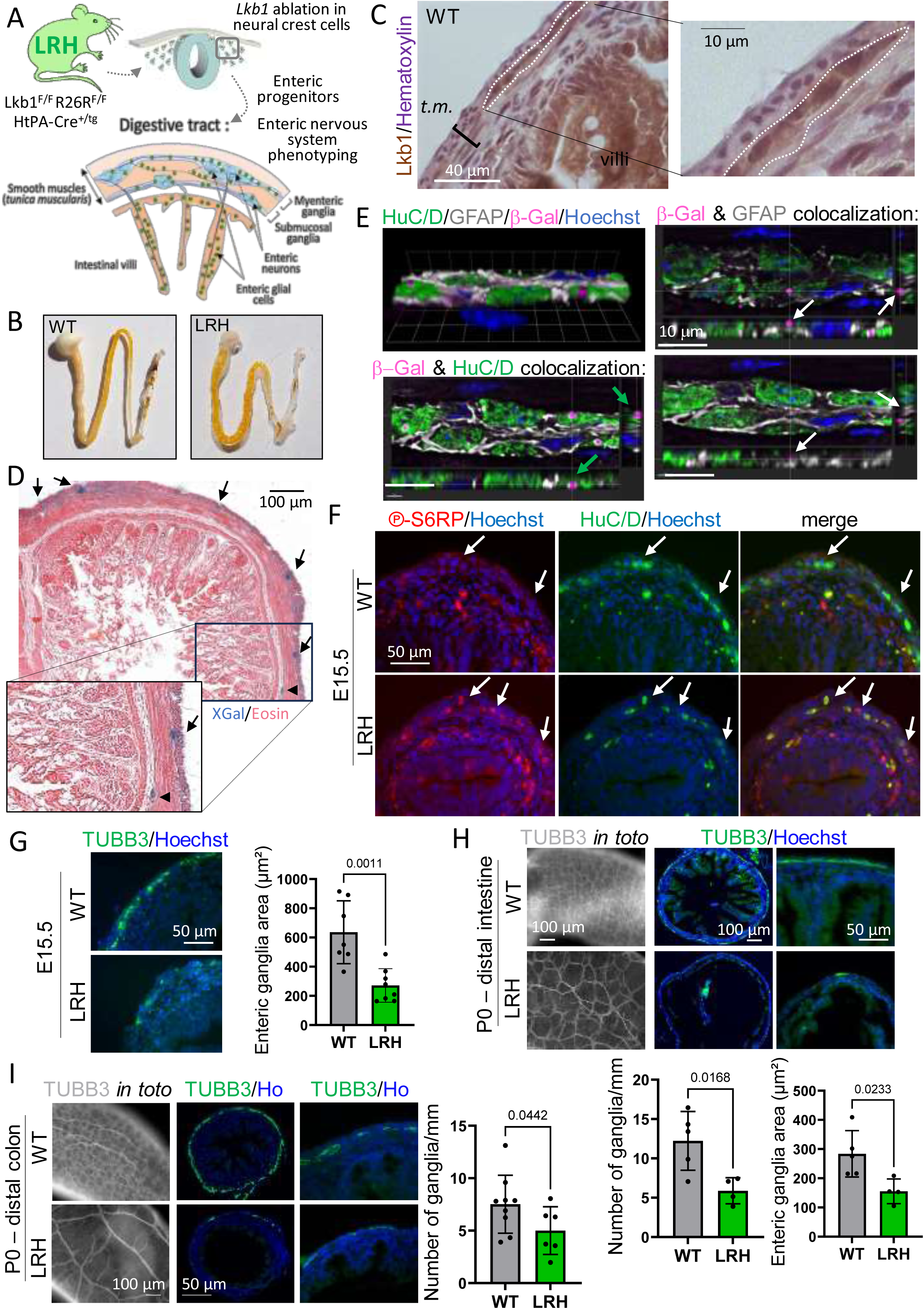
Lkb1 signaling in vagal neural crest cells is essential for enteric ganglia formation. **A**. Schematic showing spatiotemporal ablation of *Lkb1* in post-delaminating neural crest cells from E8.5, achieved by crossing mice harboring *Lkb1* floxed gene and the *LacZ* reporter **R**26R (expressing β-Galactosidase) with ***H****tPA-Cre* transgenic mice. The resulting mutants, named “LRH”, were compared to wild type (WT) animals. The diagram also illustrates the migration of enteric progenitors toward gut territories, where they form two ganglionic plexuses within the smooth muscle layers (*tunica muscularis*) and innervate intestinal villi or colonic epithelial cells. **B**. Representative images of digestive tracts from newborn animals (P0) showing air bubbles in the LRH mutants, absence of fecal material in the colon, and a thinner digestive tract compared to WT. **C**. Immunostaining of Lkb1 in P0 distal intestine transverse sections of WT mice, highlighting Lkb1 expression in enteric ganglia. **D**. X-Gal (blue) and Eosin (pink) staining of enteric ganglia (arrows) in a transverse section of the distal intestine from an adult WT mouse (Lkb1^+/+^ HtPA-Cre^+/tg^). **E**. Co-immunostaining of neurons (HuC/D), glial cells (GFAP), and β-Galactosidase (β-Gal) in enteric ganglia of a WT mouse (*Lkb1^+/+^ HtPA-Cre^+/tg^*) at P0. Confocal analysis shows Cre recombination in both enteric neurons and glial cells. **F**. Immunostaining for phosphorylated S6 ribosomal protein (p-S6RP, ser235/236) on transverse sections of the distal intestine at embryonic day 15.5 (E15.5). Increased p-S6RP staining in LRH mutants compared to WT ganglia (white arrows). WT n= 6, LRH n=5. **G**. Immunostaining of enteric neurons (TUBB3) at E15.5 in distal intestine sections of WT and LRH mice. The LRH mouse shows marked hypoganglionosis. WT n=7, LRH n=8. **H**. Visualization of the enteric network using β3-βTubulin staining on distal intestine sections (wholemount on the left, transverse sections with zoom on the right) of LRH and WT mice at P0. Quantifications of ganglia density and area (graphs below) show a nearly absent neuronal network in LRH mice, with hypoganglionic enteric nervous system (ENS). WT n=5, LRH n=4. **I**. Neuronal network staining using TUBB3 on wholemount (left) and transverse sections (middle, zoom on right images) of the distal colon at P0 in LRH and WT mice. The LRH ENS is hypoganglionic (graph on the right), and nerve fibers in LRH mutants (left bottom image) are primarily from extrinsic nerves visible due to the reduced enteric network. WT n=9, LRH n=6. Statistical significance (G-I) was determined by unpaired two-tailed t-tests and p-values are indicated. Nuclei were stained using Hematoxylin in C and Hoechst in E-I.

First, we confirmed the presence of Lkb1 protein in wildtype cells of the enteric ganglia at birth (P0) through immunohistochemical staining of transverse sections of the distal intestine (Figure 1C). Consistent with previous studies (Baas et al. 2004), *Lkb1* was also found in enterocytes within the intestinal villi. These findings are consistent with our earlier observations of LKB1 expression in mouse and human enteric ganglia (Radu et al. 2019). Additionally, the presence of β-Galactosidase, encoded by the *LacZ* reporter, was restricted to the enteric ganglia (P0, Figure 1D), similarly as previously described for the *HtPA-Cre* strain (Pietri et al. 2003). Using confocal microscopy and 3D reconstruction, we observed β-Galactosidase staining in both enteric neurons and glial cells (Figure 1E, suppl. data movie1).

LKB1 phosphorylates various downstream kinase substrates, including AMPK, which acts as an energy sensor activated under low-energy conditions (Bourouh and Marignani 2022). Once activated, AMPK initiates catabolic pathways to replenish ATP while inhibiting ATP-consuming anabolic processes, such as the mTOR protein synthesis pathway. To assess mTOR activity in enteric ganglia as a downstream indicator of LKB1/AMPK signaling, we evaluated mTOR signaling through the phosphorylation of the ribosomal protein S6RP (p-S6RP). At E15.5, we observed elevated levels of p-S6RP, suggesting reduced Lkb1/AMPK activity in enteric ganglia (Figure 1F). Overall, these findings confirm that the conditional ablation of *Lkb1* is restricted to neural crest-derived cells in the enteric nervous system and is associated with a classic loss of function in LKB1 signaling.

In this context, we observed a significant impact of *Lkb1* loss on the formation of enteric ganglia. Immunolabeling of neuronal cytoplasmic β-3 tubulin (TUBB3), a marker for enteric ganglia and their axonal projections, revealed pronounced hypoganglionosis in the distal intestine at E15.5 (Figure 1G). Whole-mount and section staining of TUBB3 demonstrated sustained hypoganglionosis due *Lkb1* loss at P0 in both the distal intestine and distal colon, which was quantified by assessing the number of enteric ganglia and their surface areas (Figures 1H,I and S1A).

In conclusion, our data establish that Lkb1 is essential for enteric gangliogenesis, and its loss in enteric progenitors leads to widespread and severe hypoganglionosis throughout the intestine and colon, indicative of impaired digestive motricity.

### Enteric Lkb1 is required for neuronal differentiation and glial cell maintenance

To examine the impact of LKB1 loss on ENS formation at the cellular level, we first quantified enteric neurons using the RNA-binding protein HuC/D, a nuclear marker for neurons. Immunofluorescent staining of HuC/D revealed a significant reduction in the number of neurons in enteric ganglia at E15.5 (Figure 2A) and in both myenteric and submucosal ganglia at P0 (Figure 2B). A decreased number of enteric neurons was also observed in the distal colon (Figure 2C).

**Figure 2.**
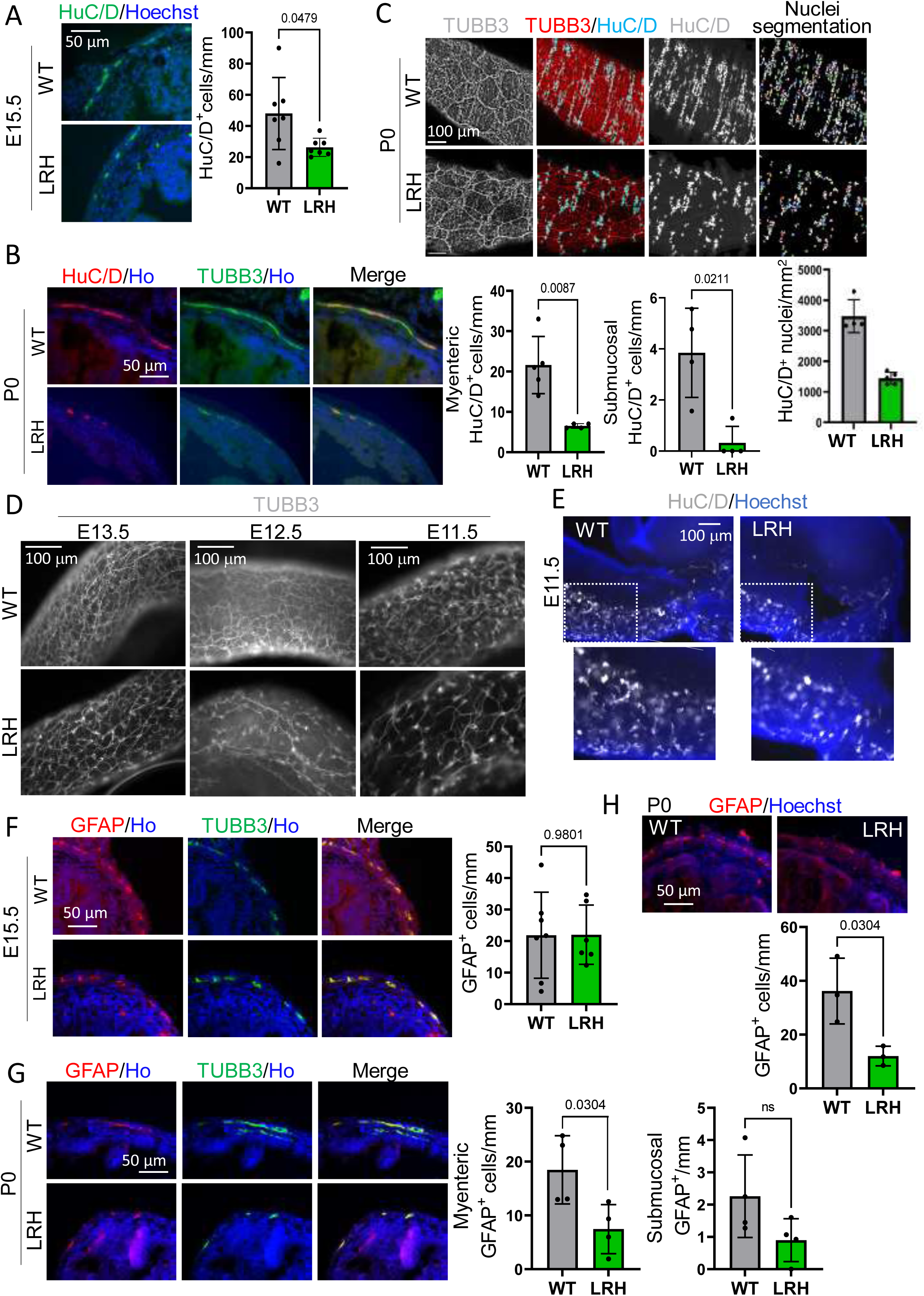
Lkb1 promotes enteric gangliogenesis through neuronal differentiation and glial maintenance. **A**. Visualization of neuronal nuclei using HuC/D labeling on transverse sections of distal intestines at E15.5. Quantification of neuron density shows a significant reduction in LRH mutants compared to WT. WT and LRH n=7. **B**. Co-immunofluorescence of HuC/D and TUBB3 on transverse sections of distal intestines at P0. A marked loss of neurons is observed in the distal intestine of LHR mice compared to WT. WT and LRH n=4. **C**. HuC/D labeling on transverse sections of the distal colon at P0. A significant loss of neurons is observed in the distal colon of LHR mouse compared to WT. WT n=4, LRH=6. **D**. Whole mount TUBB3 labeling of distal intestines at E11.5, E12.5, and E13.5. Neuronal network density is reduced in LRH mutants compared to WT at all stages. E11.5 WT n=8, cKO n=4; E12.5 WT n=7, cKO n=8; E13.5 WT n=5, cKO n=5. **E**. Wholemount HuC/D immunostaining of distal intestines at E11.5 shows a reduced number of enteric neurons in LRH mutants. Insets (below) show zoomed views. WT n=4, cKO n=5. **F**. Co-immunofluorescence of glial (GFAP) and neuronal (TUBB3) markers on transverse sections of the distal intestine at E15.5. Quantifications of GFP-positive glial cells (graph) shows no significant difference between the WT and LRH groups. WT n=7, LRH n=6. **G**. GFAP and TUBB3 labeling on transverse sections of the distal intestine at P0. There is a marked loss of both myenteric (graph on the left) and submucosal (graph on the right) glial cells in LRH mice compared to WT. WT and LRH n=4. **H**. GFAP immunostaining on transverse sections of the distal colon at P0 shows a reduced number of glial cells in LRH mutants compared to WT (graph below). WT and LRH n=3. Nuclei were stained using Hoechst. p-values of unpaired two-tailed t-tests are indicated (B, C, F-H). ns: non-significant.

To determine whether the reduced number of neurons following Lkb1 loss resulted from impaired differentiation or defective neuronal maintenance, we performed whole-mount TUBB3 immunostaining at various stages of gut colonization by NCCs (Kang et al. 2021). At all stages examined (E11.5, E12.5, and E13.5), we observed a decrease in the density of enteric neurons in the distal intestine (Figure 2D). Further analysis using HuC/D staining confirmed this reduction at E11.5 (Figure 2E). Given that enteric neurons begin to differentiate around E10 (Pham et al. 1991; Anderson et al. 2006), these results suggest that neuronal differentiation is impaired upon Lkb1 loss.

Interestingly, the number of enteric glial cells in Lkb1 mutants remained unchanged at E15.5, as indicated by GFAP staining, which marks the intermediary filaments of glial cell cytoskeleton (Figure 2F). However, later analysis at P0 revealed a massive loss of enteric glial cells in Lkb1 mutant animals, both in the distal intestine (Figure 2G) and colon (Figure 2H).

Collectively, our findings reveal a sequential role of Lkb1 in neuronal differentiation followed by the maintenance of glial cells.

### Lkb1 is not necessary for the migration or maintenance of enteric progenitors

To investigate whether *Lkb1* ablation affected the migration of enteric progenitors, we first assessed whether enteric neurons formed at the correct rostro-caudal levels. Using TUBB3 immunostaining on whole mount digestive tracts at E11.5 and E12.5, we focused on TUBB3-positive cells located closest to the migration front. Our results showed that enteric neurons in *Lkb1* null mutants successfully reached the cecum (Figure 3A, E11.5) and colonized the proximal colon (Figure S2A, E12.5) at the same developmental timing as wildtype animals. These findings suggest that *Lkb1* loss did not delay the migration of enteric progenitors.

**Figure 3.**
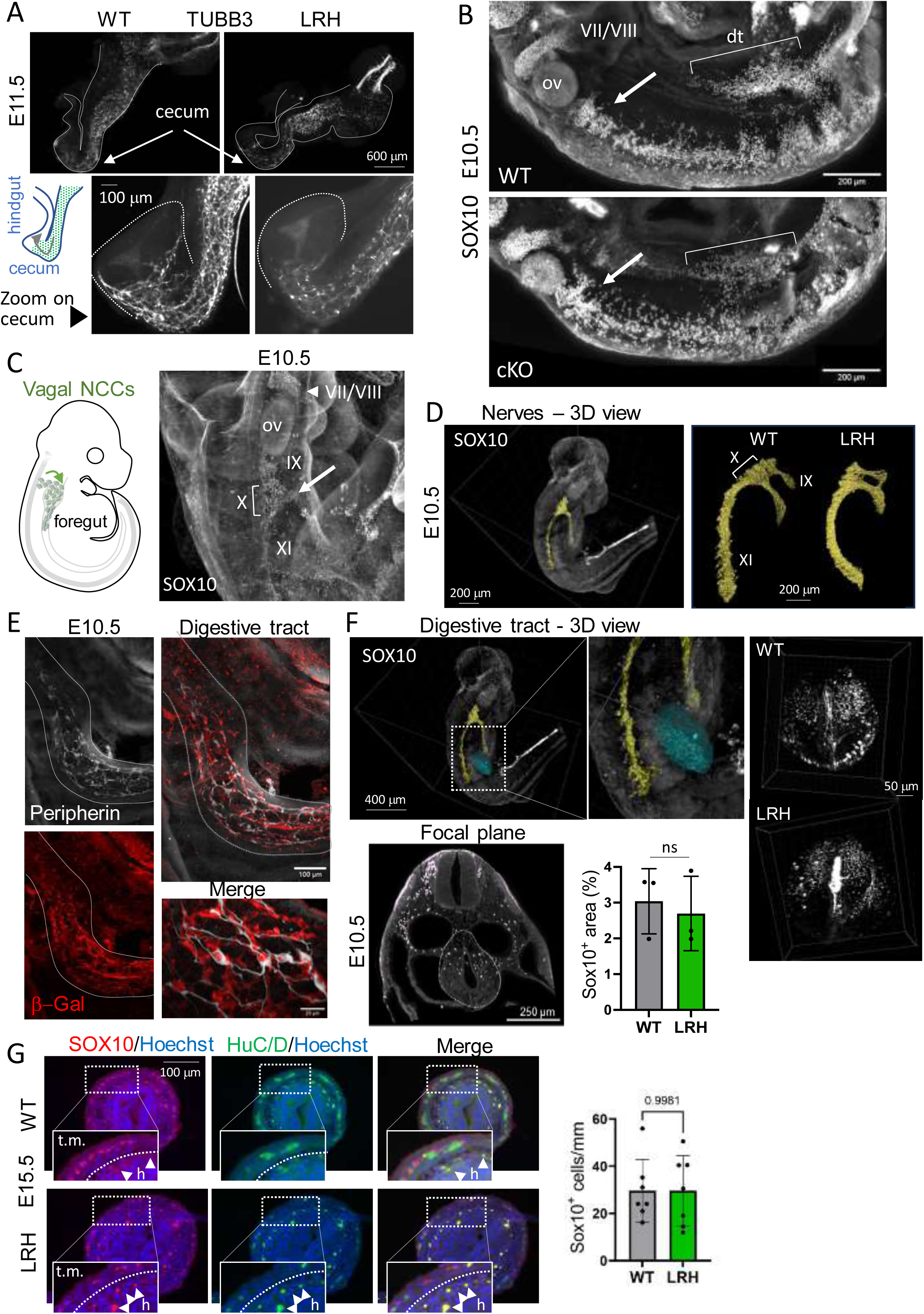
Lkb1 is not required for enteric progenitor migration nor maintenance. **A.** Wholemount TUBB3 immunolabeling of the digestive tracts (dashed lines) of LRH and WT mice at E11.5, at which stage enteric progenitors have colonized the cecum (schema on the left). Zoomed-in views of the cecum (white arrows) show TUBB3 staining, indicating that neurons are present in both groups, suggesting that *Lkb1* ablation does not impair the migration of enteric progenitors. WT n=8, cKO n=4. **B.** Side view of Maximum Intensity Projections (MIP) from cleared embryos (E10.5) after deconvolution of lightsheet microscopy images from SOX10 staining (thickness β 200 μm). White arrows indicate SOX10-positive enteric cells migrating along the vagal nerve toward the gut. Brackets show digestive tract positioning; dt: digestive tract; ov: otic vesicle. **C**. Migration path schematic (left) of enteric progenitors from the neural tube to the foregut at E10.5. 3D visualization of SOX10-positive enteric progenitors migrating along the vagal nerve (X, arrow) in cleared embryos. Cranial nerves VII (facial) and VIII (vestibuloacoustic) (arrow head), and nerves IX (glossopharyngeal) and XI (accessory) are indicated. ov: otic vesicle. **D**. Side view MIP of cleared E10.5 embryos after deconvolution of lightsheet microscopy images, showing SOX10-positive cells along nerve tracks (yellow segmentation). Segmented of nerve tracks in LRH mutants did not reveal a significant reduction in SOX10-positive cells compared to WT, but migration path defects were observed. **E**. Adaptive optics confocal imaging of Peripherin and β-Galactosidase (β-Gal) co-staining in cleared E10.5 embryos to visualize peripheral nerves. The Cre-dependent β-Gal reporter confirms *Lkb1* ablation in enteric progenitors. Co-staining shows strong overlap, as seen in zoomed images. **F**. 3D lightsheet image reconstruction of SOX10 staining showing enteric cell distribution in the digestive tract at E10.5. The white line indicates a section used for SOX10-positive population analysis (cyan segmentation). Representative image shows SOX10-positive cells in different locations within the cleared embryo. Quantification of the area occupied by SOX10-positive cells in the digestive tract shows no significant difference (Mann Whitney test) between LRH mutants and WT. WT and LRH n=3. **G**. Immunostaining of SOX10 on transverse sections of the distal intestine at E15.5 and quantification of SOX10-positive cell density. WT and LRH n=7. No statistical difference was observed between the two groups using unpaired two-tailed t-test.

Next, we aimed to investigate the impact of *Lkb1* loss on the number and migration of enteric progenitors at earlier time points. Conducting traditional histology of digestive tract at these early stages is challenging due to the organ’s small size. To overcome this limitation, lightsheet microscopy on optically cleared mouse embryo was used to visualize the positioning of enteric neurons and progenitors within the embryo (Figure S2B). We first verified the regions targeted by the recombinase Cre, driven by the *HtPA* promoter, by comparing β-Galactosidase activity from the *Cre* reporter gene (R26R) with staining of peripherin, a type III intermediate filament protein expressed in neurons of the peripheral nerves. Consistent with previous report (Pietri et al. 2003), we observed robust reporter staining in the facial region of the embryos and in components of the peripheral nervous system, such as peripheral nerves and dorsal root ganglia (Figures 3B and S3C,D). We then focused on the sympatho-enteric NCCs that emerge from the neural tube between somites 3-7 and migrate toward the gut (Espinosa-Medina et al., 2017). To identify this specific NCC subpopulation, we immunodetected the transcription factor SOX10, a well-established marker for migratory enteric progenitors (Bondurand and Sham 2013) (Figure 3C). However, we found no significant differences between wildtype and *Lkb1* mutant animals.

In addition to the sympatho-enteric subpopulation, Schwann cell precursors (SCPs), which cover the vagal nerve, also contribute to the formation of enteric ganglia (Espinosa-Medina et al. 2017; Pawolski and Schmidt 2020; Seguella and Gulbransen 2021; Lefèvre et al. 2024). To obtain a more comprehensive understanding of the entire enteric progenitor population, we assessed the morphology of nerves containing SCPs, with a focus on the vagus nerve (X), using an image segmentation approach based on SOX10 staining. No major differences were observed between wildtype and mutant animals, except for shorter branching between nerves IX (glossopharyngeal nerve) and X in *Lkb1*-ablated animals (Figure 3D).

Next, we compared the pool of enteric NCCs that were already present in the digestive tract at E10.5. To observe this population, we focused specifically on the digestive tract within embryos, identified through peripherin/βGalactosidase co-staining (Figures 3E and S2E). Confocal images with an adaptive optics confocal microscope, equipped with a long-distance, high-resolution water immersion objective, allowed us to identify cells that were positive for both markers, indicating neurons (Figure 3E and suppl. data movie2). Additionally, some cells expressed only βGalactosidase, likely representing enteric progenitors. Once again, we observed no significant differences following *Lkb1* inactivation when labeling this enteric population with SOX10, as demonstrated by quantifying the volumes of SOX10-positive cells through digestive tract segmentation (Figure 3F and suppl. data movie3). Consistent with these findings, conventional histology for SOX10 on intestine and colon sections confirmed that *Lkb1* loss did not significantly reduce the pool of SOX10-positive cells at E15.5 (Figure 3G).

Taken together, our findings indicate that, beyond the expected timeframe for *Lkb1* inactivation using the *HtPA-Cre* gene (which occurs around E8.5), Lkb1 is neither necessary for the migration nor the maintenance of enteric progenitors.

### Loss of Lkb1 in enteric cells disrupts intestinal tissue integrity and metabolism

During necropsy of LRH mutant pups, we observed that their digestive tissues appeared unusually thin and fragile (Figure 1B). Histological analyses of intestinal and colonic sections at birth revealed that *Lkb1* deletion in enteric NCCs led to a reduction in both the number and size of intestinal villi and colonic crypts, along with abnormal tissue architecture (Figure 4A). Despite this, expression of E-Cadherin, a key component of *adherens*, tight junctions and desmosomes, was largely preserved, suggesting that the intestinal epithelial barrier was not severely compromised (Figure S4A). However, other structural defects were evident, including a significant reduction in *tunica muscularis* thickness (Figure 4A), and aberrant hyperactivation of mTOR activity, as indicated by increased phosphorylation of S6RP (Figure 4B). Given LKB1’s established role in cellular metabolism, these findings suggest that its loss may disrupt metabolic homeostasis in the developing gut, thereby impairing tissue architecture and function. This prompted us to further investigate metabolic alterations in the digestive tract following *Lkb1* inactivation.

**Figure 4.**
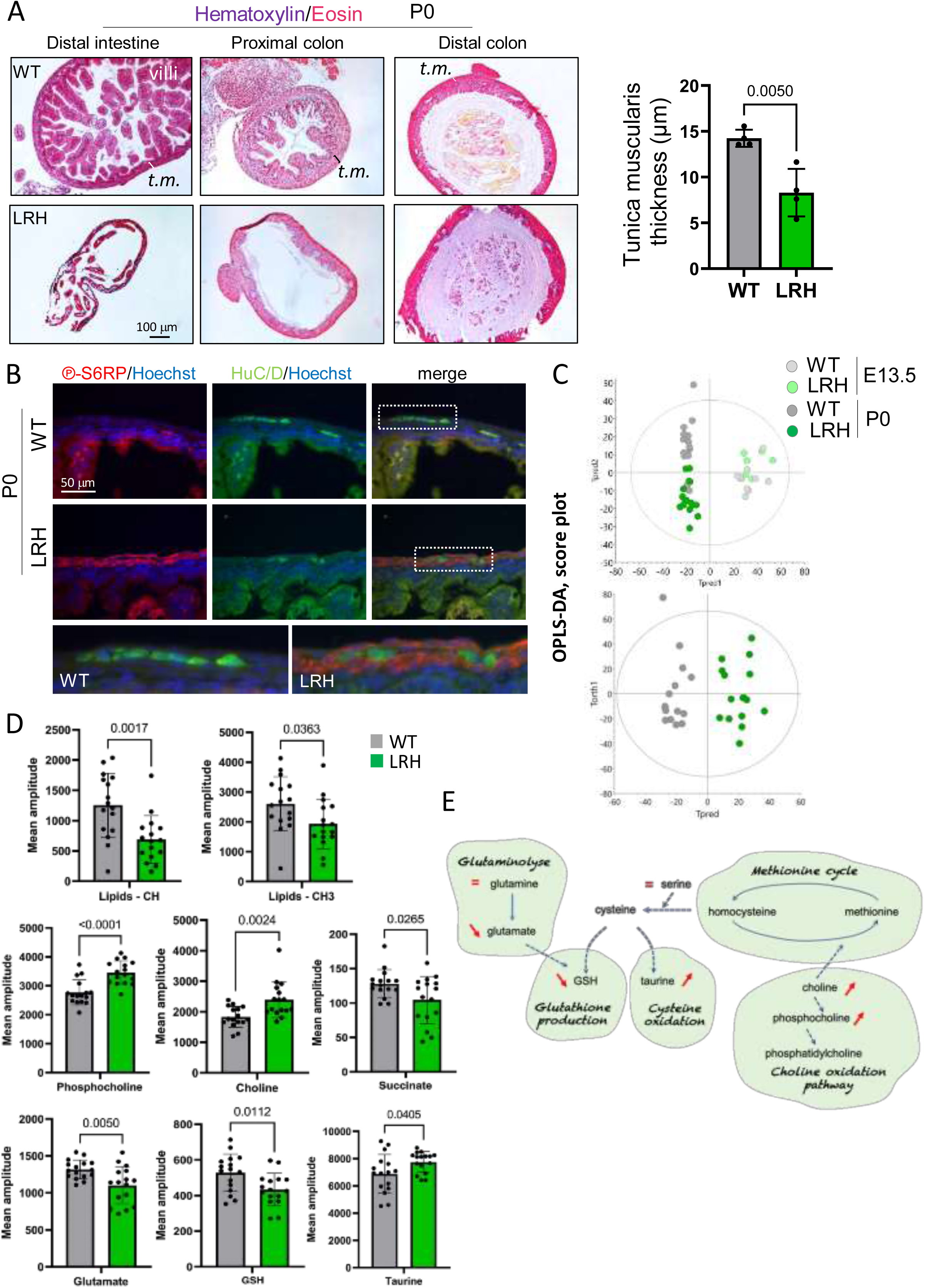
Lkb1 loss in enteric cells impairs intestinal tissue integrity and metabolism. **A**. Hematoxylin/Eosin staining of transverse sections from distal intestine, distal colon and proximal colon of P0 animals. The graph quantifies the reduction in *tunica muscularis* thickness (*t.m*., brackets). WT and LRH n=4. p-value of two-tailed unpaired t-test is shown. **B**. Immunostaining of phosphorylated S6RP on transverse sections of the distal intestine at P0. Increased S6RP phosphorylation is observed in the *tunica muscularis* of LRH mutants compared to WT, but not in enteric ganglia. WT n= 5, LRH n=4. **C.** Score scatter plot of multivariate statistical models (OPLS-DA) built from NMR spectra. Top: model incorporating both P0 and E13.5 data; Bottom: model for P0 with R^2^Y=0.902, Q^2^=0.67, and 1 predictive and 2 orthogonal components. E13.5: WT n=10, LRH n=7; P0: WT n=15, LRH n=16. **D.** Mean relative amplitude of lipids-CH and lipids-CH_3_ groups, choline, phosphocholine, succinate, glutamate, glutathione (GSH), and taurine. All chemical shifts (ο) are referenced to alanine. p-values of two-tailed unpaired t-tests are indicated. **E**. Schematic representation of metabolic pathways illustrating defects in the digestive tracts of LRH mutants.

To do so, we conducted an unbiased comparative metabolomic profiling using high-resolution magic angle spinning (HRMAS) proton-based nuclear magnetic resonance (^1^H NMR) on dissected and structurally preserved guts from E13.5 embryos and newborns (Figure S4B). We quantified 20 metabolites at P0 (Figure S4B), while only 14 metabolites were detected at E13.5 due to a much lower signal-to-noise ratio (Figure S4B). The OPLS-DA model built with the NMR data clearly identified a set of metabolites that discriminated LRH groups from WT controls at P0 (Figure 4C), all of which were significantly dysregulated (Figure S4C). In contrast, no significant differences were observed at E13.5 (Figures 4C and S4E). More specifically, *Lkb1* loss at birth was associated with a downregulation of succinate, glutamate, reduced glutathione (GSH), and fatty acid groups (CH and CH3), while phosphocholine, choline, and taurine levels were increased (Figure 4D). Interestingly, several of these metabolites are closely linked to oxidative stress adaptation (Figure 4F).

Our results indicate that *Lkb1* ablation in NCCs from E8.5 onwards progressively alters the metabolism of the entire gut between E13.5 and P0, compromising overall tissue integrity and suggesting deficiencies in redox homeostasis.

### Enteric Lkb1 regulates redox homeostasis across the entire digestive tract

Lkb1 is crucial for regulating oxidative stress (Xu et al. 2015; Tan et al. 2023), prompting us to investigate whether *Lkb1* loss leads to increased oxidative stress in enteric NCCs. To evaluate oxidative stress *in vivo*, we quantified carbonylation in total protein extracts from the digestive tracts of P0 LRH and WT mice using immunoblotting (Figure 5A). Carbonylation, induced by reactive oxygen species (ROS), serves as an early indicator of oxidative stress (Suzuki et al. 2010). In 4 out of 6 LRH mutants analyzed, we observed a significant increase in oxidized proteins, approximately twofold higher compared to the WT group.

**Figure 5.**
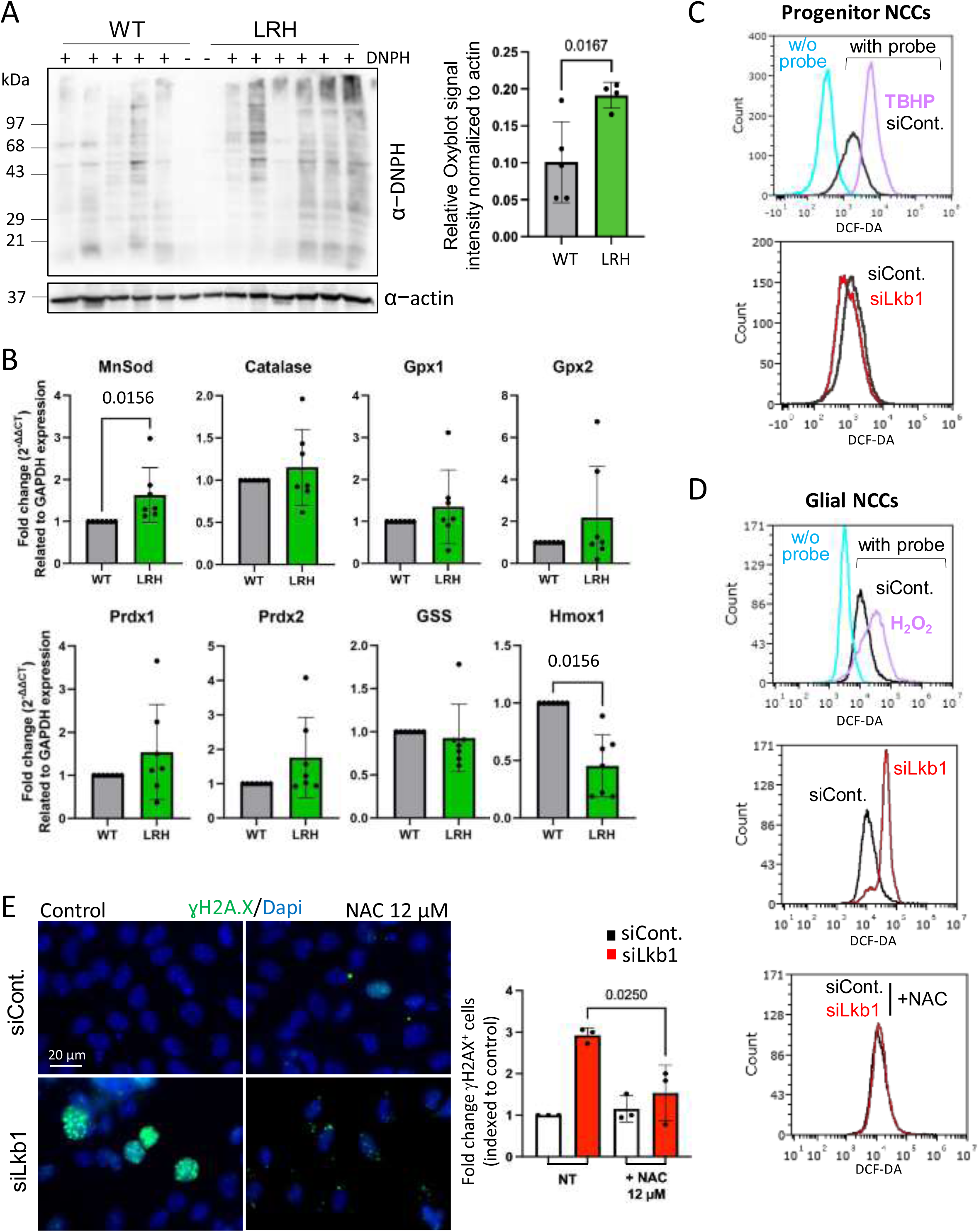
Enteric LKB1 regulates redox homeostasis in digestive tissue. **A**. Detection of oxidized protein following carbonyl derivatization with 2,4-dinitrophenylhydrazine (+ DNPH), followed by antibody detection. Protein lysates from 4 out of 6 LRH animals showed an increased signal compared to WT animals. No signal was detected in non-derivatized samples (-DNPH). Quantification was performed on 5 WT and 4 LRH mutants (graph). Protein oxidation signal was normalized to actin as a control of protein loading. p-value of two-tailed unpaired t-test is shown. **B**. Expression levels of several genes involved in oxidative homeostasis were quantified by quantitative PCR using total mRNA extracted from the digestive tracts of WT and LRH individuals. Statistically significant changes were observed in the expression of superoxide dismutase 2 (MnSOD, also known as SOD2) and heme oxygenase 1 (Hmox1) upon *Lkb1* ablation in enteric cells. Additional genes assessed include *Catalase*, *Gpx1/2* (glutathione peroxidase 1/2), *Prdx1/2* (peroxiredoxin 1/2), and *GSS* (glutathione synthetase). WT and LRH n=7, each assessed in triplicate. p-values of one-sample Wilcoxon signed-rank tests are provided where statistical differences were observed. **C**. Measurement of oxidative stress in progenitor JoMa1.3 cells *in vitro* using the dichlorodihydrofluorescein diacetate (DCF-DA) fluorescent probe to detect reactive oxygen species (ROS). Upper graph: fluorescence intensity of cells without probe (cyan), with probe treated with positive control (TBHP, pink), or with non-targeting siRNA for the control condition (siCont., black). Lower graph: representative peaks for cells treated with siCont. (black) or Lkb1-targeting siRNA (siLkb1, red). n=4. **D**. Measurements of ROS in glial JoMa1.3 cells *in vitro* using DCF-DA probe as in C. Upper graph: positive control using H_2_O_2_ (pink). Middle graph: increased ROS following *Lkb1* knockdown. Lower graph: rescue of oxidative stress following treatment with 12 μM of the anti-oxidant N-acetyl-cysteine (NAC). n=3. **E**. DNA damage labeling (“H2AX) in glial JoMa1.3 cells upon NAC treatment, showing reduced DNA damage upon *Lkb1* knockdown compared to non-treated cells (NT). The index of “H2AX-positive cells was normalized to control condition (graph). n=3. The p-value of unpaired two-tailed t-test is shown (homoscedastic variances). Total number of cells counted: siCont./NT n= 421, siCont./NAC n=295, siLkb1/NT n=187, siLkb1/NAC n=226.

We next examined the expression of oxidative stress-related genes using real-time quantitative PCR (RT-qPCR) on mRNA extracted from the digestive tract. We found a significant increase in *manganese superoxide dismutase* (*MnSOD*) expression in LRH mutants (Figure 5B). In contrast, expression levels of *glutathione peroxidase* (*Gpx1/2*), *peroxiredoxin* (*Prdx1/2*), and *catalase* genes showed only a tendency to increase, likely due to high individual variability (Figure 5C). Interestingly, the expression of the *GSS* gene, encoding glutathione synthetase, was unchanged between WT and LRH mutants (Figure S5A). On the other hand, *Heme oxygenase 1* (*Hmox1),* which encodes HO-1, was significantly reduced in LRH mutants (Figure 5D).

To determine whether ROS in the digestive tract originates from NCCs, we used the JoMa1.3 *in vitro* model of neural crest stem cells, which can be maintained as progenitors or differentiated into various neural crest derivatives (Maurer et al. 2007; Radu et al. 2019). We had previously shown that *Lkb1* knockdown impairs glial cell specification from NCCs via disrupted pyruvate-alanine conversion and altered mTOR activity (Radu et al. 2019). While adhesion and survival of progenitor NCCs were also impacted by *Lkb1* knockdown, alanine levels remained unaffected in this population. We thus investigated the role of Lkb1 in regulating oxidative stress in progenitor and glial differentiated NCCs. Using the DCF-DA fluorogenic probe to detect intracellular ROS, we found that *Lkb1* knockdown was associated with elevated ROS levels in glial NCCs but not in progenitors, compared to control siRNA-transfected JoMa1.3 cells (Figures 5E and F; Figure S5A). In progenitor NCCs, the expression of oxidative stress-related genes was not drastically deregulated by *Lkb1* knockdown, except for *Gpx1* and *Gpx2* genes which were down– and up-regulated respectively (Figure S5B). Collectively, these results suggest that Lkb1 regulates redox homeostasis at multiple cellular levels, operating through complex and context-dependent mechanisms.

Since ROS can induce DNA damage (Shadfar et al. 2023) and LKB1 preserves genome integrity through the BRCA1-mediated DNA repair pathway (Gupta et al. 2015), we assessed DNA damage in JoMa1.3 glial cells using immunofluorescence staining for phosphorylated histone H2A.X (γH2A.X), a key marker of DNA double strand breaks, especially in response to oxidative stress. *Lkb1* knockdown in glial cells led to significantly increased γH2A.X staining, indicating unrepaired DNA damage (Figure S5C). Interestingly, this DNA damage was alleviated by treatment with the antioxidant N-acetylcysteine (NAC), as evidence by reduced γH2A.X staining in *Lkb1* knockdown glial cells (Figure 5G).

We next investigated whether antioxidant treatment could mitigate the enteric phenotype in LRH mutants. NAC (0.4g/kg/day) was administered via drinking water to pregnant and lactating females (Figure S5D), but this did not rescue the neurocristopathy phenotype or improve the survival of mutant mice. Furthermore, NAC treatment adversely affected the weight gain of wildtype neonates (Figure S5E). For both scientific and ethical reasons, we decided to discontinue further antioxidant treatment and shifted focus to investigating downstream molecular effectors.

In conclusion, our findings establish that *Lkb1* ablation in NCCs leads to the accumulation of oxidative stress throughout the entire digestive tract, not only in glial enteric cells but also in adjacent digestive tract tissues. This suggests that Lkb1 plays a crucial role in mediating redox homeostasis and mediating communication between enteric cells and surrounding mesenchymal and epithelial tissues, including smooth muscle and intestinal villi cells.

### Lkb1 loss activates p53 and induces apoptotic cell death in enteric ganglia

To identify the downstream effectors of Lkb1 loss, we performed transcriptomic profiling (RNAseq) of JoMa1.3 progenitors following *Lkb1* knockdown. Among the differentially expressed genes (DEGs), 18 were significantly downregulated, and 11 were upregulated (Figure 6A). As expected, *Lkb1* expression was reduced (Fig S5F), and we also observed changes in several genes involved in metabolic processes (*e.g., Mat2a*, *Aldh1l2*, *ATF5, Rsc1a1*), with *Chac1* and *Aldh1l2* playing key roles in redox homeostasis. Additionally, genes related to cell adhesion (*e.g., Itga6*, *Msln*, *Acan*, *Adamts17*) and neuronal development (*e.g., Penk*, *Chac1*) were also impacted. GO term analysis of DEGs revealed significant impairments in pathways related to the cell cycle, adhesion, neuronal development, metabolic processes, and autophagy (Figure 6B, S5G). Notably, we identified deregulated genes linked with apoptosis (Figure 6B), including a marked upregulation of p53 (Figure S5H).

**Figure 6.**
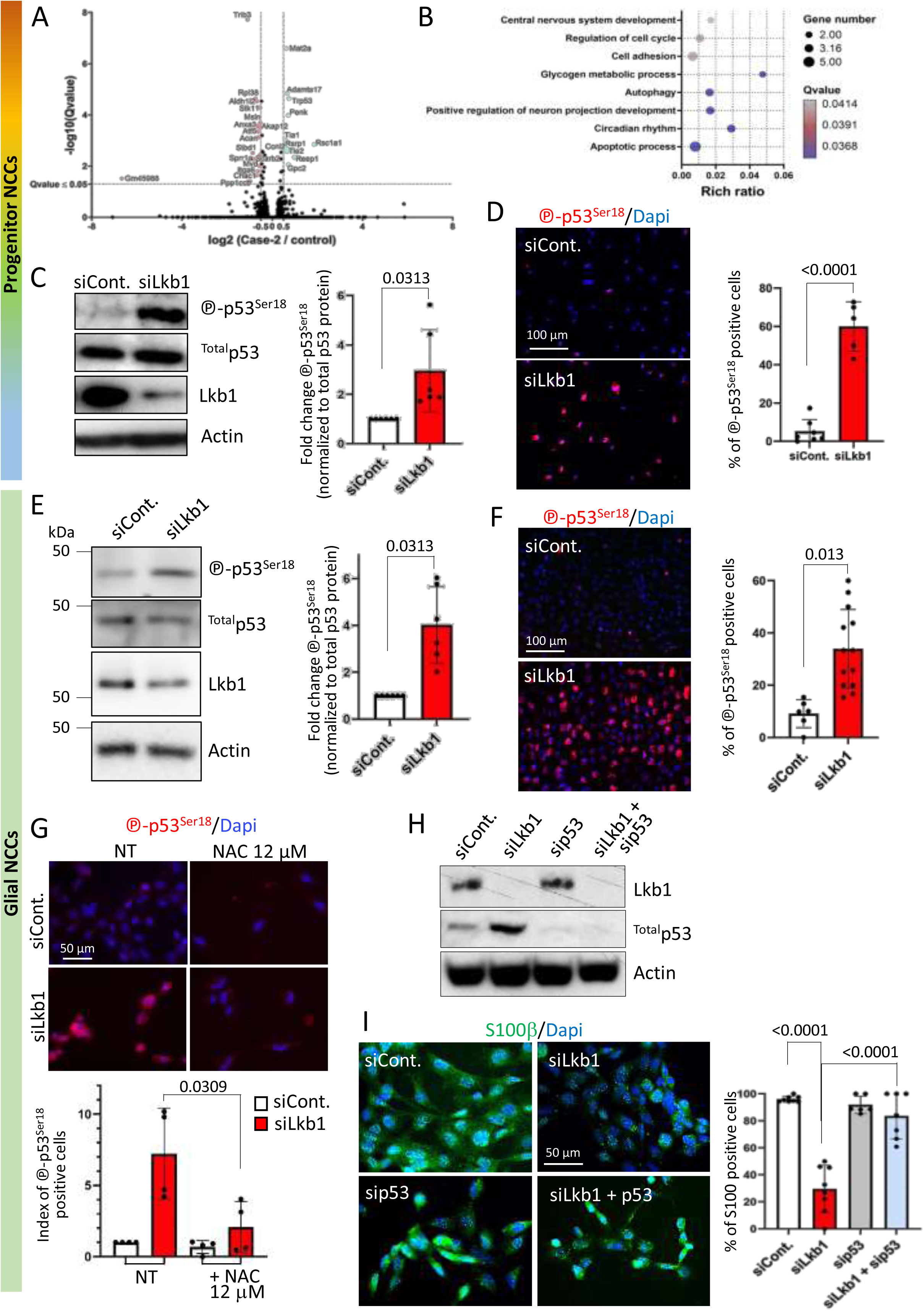
Lkb1 regulates ROS and p53 activity during glial specification and maintenance *in vitro* **A**. Volcano plot analysis showing downregulated (red) and upregulated (cyan) genes upon *Lkb1* knockdown in progenitor JoMa1.3 cells. mRNA data from cells of the two conditions were compared for each gene, providing a unique significance score and fold change. n=2. **B**. Enrichment of differentially expressed genes (both down– and up-regulated) in *Lkb1* knockdown progenitors for Gene Ontology (GO) terms related to biological processes (BP), with FDR<0.05. **C**. Western blot analysis of protein extracts from progenitor JoMa1.3 cells showing accumulation of phosphorylated p53 at Ser18 (℗-p53^Ser18^). Quantification represents the fold change of p53 phosphorylation level normalized to actin and total p53 protein. n=6. **D.** Immunolabelling of p53 phosphorylation at Ser18 (℗-p53^Ser18^) *in vitro* using the neural crest cell line JoMa1.3 maintained as progenitor cells. *Lkb1* knockdown using Lkb1-targeting siRNA (siLkb1) compared to non-targeting siRNA as control condition (siCont.) triggered increased p53 phosphorylation. Representative images of epifluorescence microscopy are shown with quantification of the immunolabeling (graph) showing significant increase of percentage of cells with phosphorylated p53 upon *Lkb1* knockdown. n=3. Total number of cells counted: siCont = 889, siLkb1 = 241. p-value of unpaired two-tailed t-test is shown. **E.** Same as in C with glial JoMa1.3 cells upon Lkb1-targeting siRNA (siLkb1). n=6. **F.** Same phospho-p53 immunofluorescence as in D performed in glial JoMa1.3 cells. n=3. Total number of cells counted: siCont. n=171, siLkb1 n=187. **G.** Accumulation of phosphorylated p53 upon *Lkb1* knockdown in glial JoMa1.3 cells is limited by the antioxidant NAC treatment compared to untreated cells (NT). Quantifications are shown (graph) n=4. p-value of t-test is indicated. Total number of counted cells: siCont. NT n=377, siCont. NAC n=167, siLkb1 NT n=339, siLkb1 NAC n=285. **H.** Western blot validation of combined *Lkb1* and *p53* knockdown by RNA interference (siLkb1+sip53) in JoMa1.3 cells differentiated toward the glial lineage. The combined application of siRNAs effectively reduced both Lkb1 and total p53 protein levels. n=2. **I**. Glial-committed JoMa1.3 cells with *Lkb1* knockdown (siLkb1) showed decreased labelling by the glial marker S100 compared to non-targeting control siRNA (top images). However, combined *Lkb1* and *p53* knockdown (siLkb1+sip53) rescued glial cell differentiation (bottom images). The percentage of S100β-positive cells is shown (graph). siCont. n=288, siLkb1 n=356, sip53 n=234, siLkb1+sip53 n=190. n=2. For Western blot analyses, statistical significance was tested using Wilcoxon signed rank test (two-tailed) and p-values are indicated. For immunofluorescence p-values of unpaired two-tailed t-test are shown (homoscedastic variances).

The transcription factor and tumor suppressor p53 is activated in response to cellular stressors, such as DNA damage (Shadfar et al. 2023). Upon activation, p53 triggers various stress-response mechanisms, including cell cycle arrest, DNA repair, and apoptosis. Misregulated p53 activation during embryogenesis can lead to developmental defects, particularly in NCC ontogenesis (Bowen and Attardi 2019; Bowen et al. 2019). To investigate whether Lkb1 loss affects p53 signaling, we measured p53 levels and activity in JoMa1.3 progenitor cells, using the phosphorylation of Ser18 in mouse p53 (equivalent to Ser15 in human p53) as a marker of p53 activation (Meek and Anderson 2009). Both western blot analyses (Figure 6D) and immunofluorescence staining (Figure 6E) revealed an accumulation of phosphorylated p53 following *Lkb1* knockdown in JoMa1.3 progenitor cells. Similar p53 stabilization was observed in JoMa1.3 cells committed to glial specification (Figures 6E, F).

Given that neurons are particularly sensitive to p53 activation (Bowen et al. 2019), we hypothesized that Lkb1 could also play a role in neuronal differentiation through the regulation of p53 activity. To test this hypothesis, we differentiated JoMa1.3 cells toward the neuronal lineage (Figures S6A-C) and observed that *Lkb1* knockdown inhibited neuronal differentiation *in vitro* (Figures S6D-F), which was associated with deregulated AMPK/mTOR activities (Figure S6G), consistent with previous findings in progenitor and glial NCCs (Radu et al. 2019). Interestingly, *Lkb1* knockdown in neurons did not affect alanine levels (Figure S6H), unlike the changes observed in glial cells. Additionally, mitochondrial topology and mass were altered by *Lkb1* knockdown (Figures S6I, J), although mitochondrial activity remained unaffected (Figure S6K). Surprisingly, p53 stabilization was not observed in neuronal derivatives from NCCs following *Lkb1* knockdown (Figure S6L), suggesting a cell-type specificity in the Lkb1-p53 balance in progenitors and glial NCCs.

To confirm that p53 activation in glial NCCs resulted from oxidative stress induced by *Lkb1* loss, we treated JoMa1.3 glial cells with NAC, which effectively prevented the accumulation of phosphorylated p53 following *Lkb1* knockdown (Figure 6G).

Genetic ablation of *p53* has been shown to prevent NCC malformations in several neurocristopathy models (Bowen and Attardi 2019). Therefore, we investigated whether *p53* ablation could rescue the NCC defects caused by Lkb1 loss *in vitro* using JoMa1.3 glial cells. Double knockdown of both *Lkb1* and *p53* was confirmed by western blot (Figure 6H). in these cells, glial differentiation and maintenance, as indicated by the S100β marker, was significantly restored compared to cells with *Lkb1* knockdown alone (Figure 6I).

Together, these results highlight the cell-type specificity of the LKB1-p53 axis and suggest that p53 activation plays a critical role in the impaired glial differentiation and maintenance observed in the absence of Lkb1.

### p53 deletion partially restores the enteric axonal network and gut tissue homeostasis

Next, we evaluated p53 activation *in vivo* in LRH mutants compared to wildtype animals *in vivo*, using lightsheet and adaptive optics confocal microscopy on cleared E10.5 mouse embryos. We initially observed an increased signal of phosphorylated p53 throughout the analyzed sections of LRH mutant embryos, indicating widespread tissue activation (Figure 7A). In parallel, we quantified an elevated phospho-p53 staining in the dorsal region of the digestive tract that overlapped with previously identified SOX10-positive cells (Figure 7A, zooms on the right and supdata movie4). Accumulation of phosphorylated p53 was also detected in enteric ganglia by conventional histological analysis of digestive tract sections from E15.5 embryos (Figure S7A), with both enteric neurons and glial cells showing positive staining (Figure S7B). Elevated p53 phosphorylation was also observed in the few remaining ganglia of LRH newborn pups (Figure S7C).

**Figure 7.**
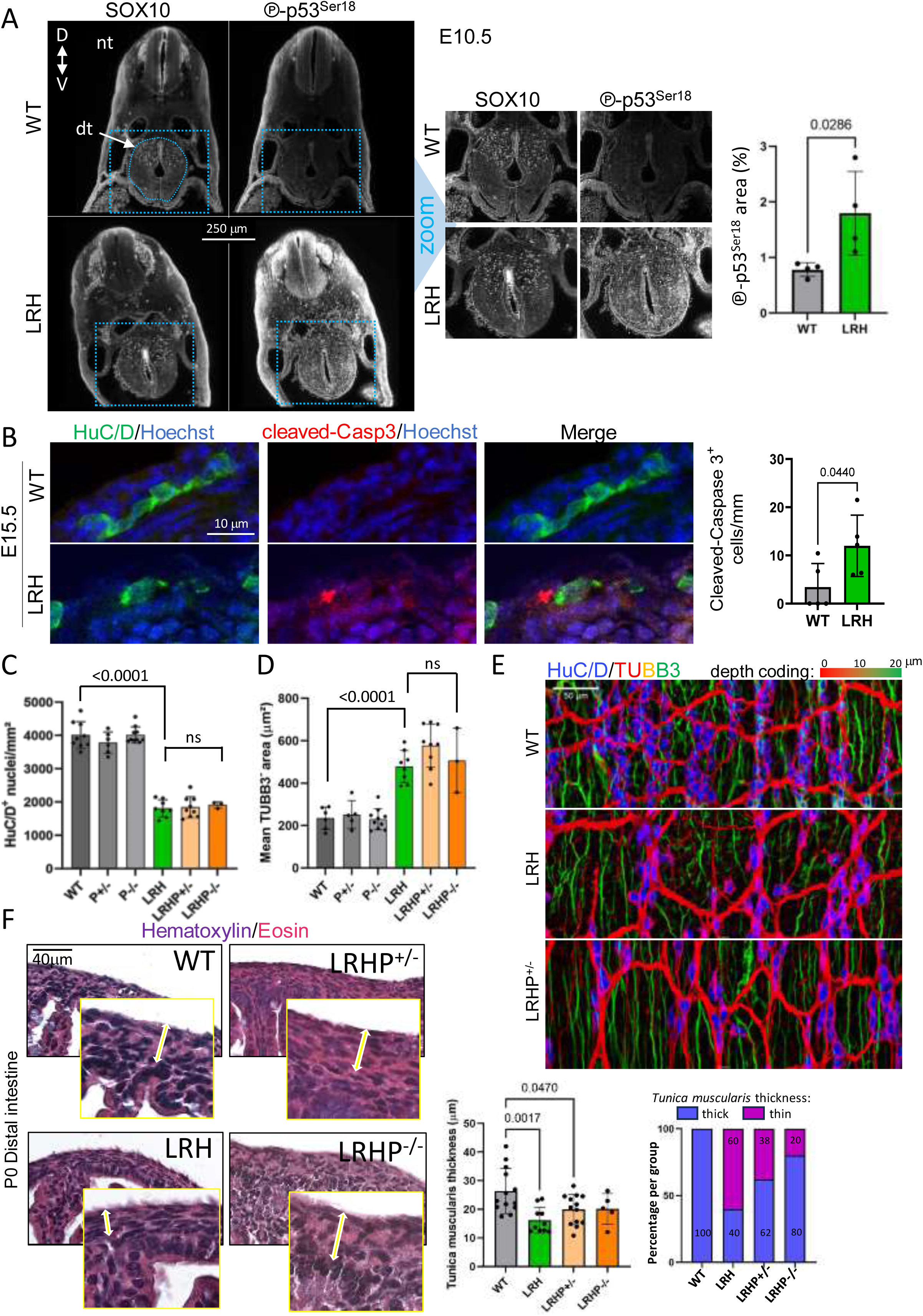
Lkb1 is essential for enteric glial cells and whole tissue homeostasis partly though p53-dependent regulatory mechanisms. **A**. 3D imaging of cleared E10.5 embryos using light sheet microscopy showing SOX10 and phosphorylated p53 (℗-p53) labeling. Inset squares with dashed lines zooming on transverse focal planes of the digestive tract are shown on the right. Acquisitions concerning the region of interest focused on the digestive tracts correspond to a stack of 51 images (100 μM) and were deconvoluted. Phosphorylated p53-positive staining was quantified and indicate a significant increase of the signal in LRH mutants compared to WT mice (graph). WT and LRH n=4. **B**. Activated caspase 3 labeling (cleaved-Casp3) on transverse sections of intestine in mice at E15.5 showing staining in the vicinity of enteric ganglia. WT and LRH n=5. p-values of two-tailed t-tests are indicated on graphs. **C**. Quantification of the density of HuC/D-positive nuclei after segmentation (graph). WT n=9, LRH n=8, LRHP^+/-^ n=9, LRHP^-/-^ n=2. Controls with wildtype *Lkb1* alleles but loss of one or two *p53* alleles are also shown (P^+/-^ n=6, P^-/-^ n=10, respectively). Statistical analyses were performed using two-tailed unpaired t-test. **D**. Quantification of TUBB3-negative areas also show no rescue of Lkb1-mediated increased TUBB3-negative areas when *p53* was deleted. WT n=9, P^+/-^ n=5, P^-/-^ n=10, LRH n=8, LRHP^+/-^ n=9, LRHP^-/-^ n=2. Statistical comparisons were conducted using two-tailed unpaired t-test. ns: non-significant. **E.** Depth-coded imaging in P0 whole-mount distal intestines using adaptive optics confocal microscopy enabled visualization of both superficial and deeper nerve fibers. **F.** Hematoxylin and Eosin staining of transverse sections from the P0 distal intestine was used to measure *tunica muscularis* thickness (center graph). The proportion of individuals with thin (<18 mm) or thick *tunica muscularis* is shown on the right. WT n=13, LRH n=10, LRHP^+/-^ n=13, LRHP^-/-^ n=5.

To investigate whether p53-driven apoptosis contributes to the loss of enteric glial cells, we assessed apoptosis *in vivo* using active caspase-3 as a marker (Bowen and Attardi 2019). Immunostaining of E15.5 digestive tract sections revealed significantly increased apoptosis in the enteric ganglia of LRH mutants compared to wildtype controls (Figure 7B). Apoptotic cells were also detectable as early as E13.5 via whole-mount staining, with higher incidence in LRH mutants (Figure S7D). In newborn pups, however, extensive tissue disorganization caused by *Lkb1* loss hindered reliable detection of caspase-3 positive cells (Figure S7E).

Taken together, these findings demonstrate that the loss of *Lkb1* in enteric NCCs leads to oxidative stress, DNA damage, p53 activation, and ultimately, apoptosis in the enteric ganglia. Thus, the Lkb1-mediated regulation of p53 signaling is essential for the proper development of the ENS.

We next investigated the impact of p53 loss *in vivo* by breeding LRH mice with *p53* knockout mice (constitutive ablation model) (Figure S8A). This breeding produced mice with *Lkb1* loss specifically in NCCs, along with the deletion of one or two *p53* alleles in all cells (LRHP^+/-^ and LRHP^-/-^ animals). Weight monitoring from postnatal day 5 until survival showed no improvement with the loss of one *p53* allele (Figure S8B).

LRHP^+/-^ animals displayed the typical complex neurocristopathy phenotype, including coat depigmentation and motor defects, such as hindlimb paralysis (Figure S8C). Obtaining LRHP^-/-^ animals was challenging, primarily due to the low expected frequency of this genotype (1/16, Figure S8A). Furthermore, although LRH mutants followed a Mendelian distribution at all embryonic stages examined (Figure S8D), their survival rate at birth was reduced to 58%, based on a total of 342 newborns (Figures S8E,F), consistent with our previous findings (Creuzet et al. 2016). Statistical analysis of Mendelian inheritance in 703 newborns revealed that the loss of one or two *p53* alleles in LRH background did not significantly alter birth survival rate (either improved or worsen) (Figure S8E).

Given the subtlety of any potential rescue of the neurocristopathy phenotype – especially considering that the survival rate of LRH mutants was not altered by p53 ablation-we conducted detailed quantification of enteric nuclei and the axonal network. Using z-stack reconstructions from adaptive optics confocal imaging of the digestive tracts of newborn pups (Figure S9), we observed no significant differences in the number of HuC/D-positive enteric nuclei (Figure 7C) or in the area of the TUBB3-positive axonal network (Figure 7D) following deletion of one or both *p53* alleles in LRH mutants. However, the organization and directionality of thin nerve fibers showed partial rescue upon p53 loss (Figure 7E). Assessment of global tissue homeostasis, based on *tunica muscularis* thickness, revealed improvement following p53 deletion, with a greater proportion of individuals displaying restored thickness in a dose-dependent manner relative to the number of p53 alleles deleted (Figure 7F).

Taken together, these results suggest that both p53-dependent and p53-independent mechanisms contribute to the neurocristopathy phenotype associated with Lkb1 loss.

## Discussion

### Multifaceted roles of Lkb1 signaling in enteric ganglia

This study demonstrates that conditional ablation of *Lkb1* in pre-migrating NCCs using the *HtPA-Cre* driver leads to severe hypoganglionosis and disrupted gut tissue homeostasis, suggesting impaired digestive motility.

Our findings reveal that Lkb1 is crucial for enteric neuron differentiation as early as E11.5, while its role in maintaining enteric glial cells appears more critical than in their differentiation.

As previously shown, Lkb1 is also essential for postnatal maintenance of enteric ganglia (Radu et al. 2019). In a genetically engineered mouse model with *Lkb1* disruption in post-migratory NCCs starting at E10.5, ENS formation was unaltered at E15.5 and birth. However, there was a reduction in enteric ganglia in the distal intestine and colon, more pronounced with age, observed at 6 and 21 days postnatally. This suggests that *Lkb1* loss during later stages of NCC development leads to progressive postnatal enteric degeneration. Combined with our findings of Lkb1 inactivation in pre-migrating NCCs, these results highlight the multifaceted regulatory role of Lkb1 signaling within the ENS. In light of previous findings regarding Lkb1’s involvement in peripheral nerves (Ulisse et al. 2020; Beirowski et al. 2014a; Radu et al. 2019; Shen et al. 2014; Pooya et al. 2014), our data firmly establish Lkb1 as a master regulator of peripheral neurons and associated glial cells.

### Differential regulation of glial cell specification and maintenance by Lkb1

While *Lkb1* loss impaired the maintenance of enteric glial cells, it did not hinder their differentiation. In contrast, Lkb1 is essential for Schwann cell differentiation and maintenance in peripheral nerves, both in *in vitro* and *in vivo* (Radu et al. 2019; Beirowski et al. 2014b; Pooya et al. 2014; Shen et al. 2014). Several factors may explain this discrepancy. First, the timing of gliogenesis in the ENS occurs earlier than in peripheral nerves, with enteric glial cell differentiation occurring around E11.5-12, after enteric neurons (Charrier and Pilon 2017), whereas Schwann cell differentiation is postnatal (Jessen and Mirsky 2019). Lkb1’s role in glial maturation may thus be more critical in the postnatal period. Additionally, neuro-glial communication, essential for glial cell maintenance may be disrupted in Lkb1 mutants, as glial maturation depends on juxtracrine signals from neurons (Newbern 2015; Espinosa-Medina et al. 2017). Furthermore, intrinsic or extrinsic factors like reactive oxygen species (ROS) from surrounding cells could contribute to glial cell death during late-stage embryogenesis. Lastly, Schwann cells have distinct lipid and protein metabolic requirements for myelination, which depend on Lkb1-mediated metabolic regulation (Beirowski 2018; Thibert et al. 2023), whereas enteric glial cells likely have different metabolic demands, though this topic remains largely unexplored.

### Metabolic insights into enteric NCC development and function

So far, our understanding of the metabolic requirements of NCC subpopulations is indeed limited. Cranial NCCs require enhanced aerobic glycolysis for delamination and migration (Bhattacharya et al. 2020), while trunk NCCs rely on glucose oxidation via oxidative phosphorylation for their prominent phenotype (Nekooie Marnany et al. 2023). Maturating Schwann cells, which produce myelin, also depend on oxidative phosphorylation, a process regulated by LKB1 (Pooya et al. 2014; Beirowski et al. 2014a). However, impaired glial specification due to Lkb1 knockdown could be rescued independently of the mitochondrial activity, only by acting on pyruvate-alanine cycling and mTOR (Radu et al. 2019). Despite multiomics-based advances in understanding the molecular mechanisms underlying enteric lineage formation (Wright et al. 2021; Zhou et al. 2024 as examples among others), the metabolic profiles of enteric progenitors, neurons and glial cells remain largely unexplored.

We employed nuclear magnetic resonance (NMR) profiling, carbonylated protein detection, and gene analysis of redox homeostasis to investigate this. Our results showed that *Lkb1* loss triggers an oxidative stress condition, with reductions in glutamate and glutathione metabolites across the digestive tracts in neonates, but not in E13.5 embryos, indicating the gradual onset of oxidative stress during development.

Additionally, we observed downregulated expression of the oxidative stress marker *Hmox1* (encoding Heme Oxygenase), a key enzyme involved in immunity and inflammation (Ryter 2022). AMPK modulates *Hmox1* transcription by regulating NRF2 either through direct phosphorylation (Matzinger et al. 2020) or by negatively regulating the NRF2 cofactor BACH1 (Fischhuber et al. 2020). Future exploration of the regulation of *Hmox1* in LRH mutants would be deserved.

Identifying the source of ROS in Lkb1 mutants is also crucial for targeting potential therapeutic interventions. One possibility is mitochondrial dysfunction resulting from impaired oxidative phosphorylation, as noted in several *Lkb1*-deficient models (Beirowski et al. 2014a; Pooya et al. 2014; Radu et al. 2019) and in the JoMa1.3 cell line *in vitro* (Radu et al. 2019). ROS accumulation *in vitro* was observed in both glial cells and progenitors, suggesting a conserved role for Lkb1 across the neural crest lineage. However, NAC treatment *in vitro* inhibited glial differentiation in control JoMa1.3 cells, even at low doses. Similarly, our *in vivo* observations revealed that NAC treatment negatively impacted the weight gain of wildtype newborn animals without improving the condition of LRH mutants. These findings align with the physiological roles of ROS, which can have distinct effects at critical thresholds, both at low and high concentrations (Sies and Jones 2020). For instance, low concentrations of H_2_O_2_ can hinder neural differentiation and maintain progenitors in quiescent state, while excessively high concentrations can trigger axonal degeneration. In this context, exploring how Lkb1 regulates the ENS through the recruitment of AMPK or other AMPK-related kinases downstream of Lkb1 would be valuable, especially since AMPK activity is influenced by redox control (Sies and Jones, 2020).

*In vitro,* we demonstrated that *Lkb1* loss leads to ROS-dependent p53 activation both in progenitors and glial-specified NCCs. However, the density of enteric progenitors was not decreased at least in early developmental stages (as assessed as E10.5 and E15.5), and only the pool of enteric glial cells was decreased from E15.5 following Lkb1 inactivation. Consequently, it is crucial to understand why enteric progenitors are resilient to ROS-p53-mediated cell death signals.

### LKB1-p53 balance in enteric neural crest cells

Increased p53 activity has been implicated in a range of developmental disorders in both human patients and mouse models (Bowen and Attardi 2019). The extent and spatiotemporal pattern of aberrant p53 activation can trigger apoptosis and cell-cycle arrest in a tissue-specific manner. Importantly, apoptotic outcomes appear to be governed more by gene expression profiles than by mitochondrial priming (Bowen et al. 2019).

Neurons derived from trunk NCCs exhibit heightened sensitiv to p53-dependent apoptosis compared to their cranial counterparts and to more resistant NCC-derived mesenchymal cells, highlighting lineage-specific variability in stress responses (Bowen et al. 2019).

In our model, loss of Lkb1 leads to p53 stabilization in enteric progenitors, which coincides with impaired neuronal differentiation. This aligns with findings from mouse models with stabilized p53, in which transcriptomic analysis revealed a significant downregulation of neuronal gene signatures (Bowen et al. 2019). These observations suggest that inappropriate activation of p53 disrupts enteric neurogenesis, likely contributing to the hypoganglionosis observed following Lkb1 loss.

However, deletion of one or both *p53* alleles did not rescue the neurocristopathy phenotype, suggesting that recovery may be hindered by the multifaceted roles of p53 during early development, particularly in regulating both differentiation and apoptosis (Raj et al. 2022), or by p53-independent pathways downstream of Lkb1.

### Implications for human disease and therapeutic applications

Mutations in *LKB1* (*STK11*) have been linked to cachexia, a wasting syndrome common in cancer patients and which causes significant muscle and adipose tissue loss (Lyengar et al. 2023). *LKB1* mutations were found to induce cachexia through immune modulation of the tumor microenvironment (Lyengar et al. 2023). Our Lkb1-deficient mouse models also exhibit weight loss and disrupted tissue homeostasis, as observed in this study and our previous work (Radu et al. 2019). Therefore, it would be valuable to explore whether cachexia-related markers are implicated in *Lkb1*-deficient tissues.

Additionally, LKB1 has long been recognized as a mediator of intercellular communication, facilitating signal exchange between different cell types, sometime over long distances, through secreted molecules like TGFβ (Ollila and Mäkelä 2011). This is reminiscent of the concept of “landscaper” genes, which encode proteins that regulate the microenvironment by modulating interactions between cells or between cells and the extracellular matrix. Consequently, LKB1’s landscaper activity within NCCs may contribute to the phenotypes observed in our genetic models of *Lkb1* inactivation.

Our findings broaden the potential involvement of defective LKB1 signaling in a spectrum of human diseases linked with gastrointestinal motility disorders, including neurodevelopmental and neurodegenerative conditions (Holland et al. 2021). Considering the promising therapeutic potential of enteric neural crest stem cells in regenerative medicine for treating neurogliopathies (Holland et al. 2021; Rahman et al. 2024), further investigation into LKB1-dependent molecular mechanisms regulating ENS biology is essential.

## Methods

### Animals Husbandry

Mouse lines were maintained at « Plateforme de Haute Technologie Animale (PHTA)» UGA core facility hTAG, Inserm US46, CNRS URA2019 (La Tronche, France, EU0197, Agreement D38-516 10 006). Animals for study cohorts were generated at the Institute for Advanced Biosciences (IAB, Agreement D38-516 10 001). Animals were housed in both facilities in accordance with European communities Council Directive 2010/63/EU, under specific pathogen–free conditions, temperature-controlled environment with a 12-h light/dark cycle and *ad libitum* access to water and diet. Protocols involving animals were reviewed by the local ethic committee (CEEA #012) and approved by the French Ministry of Higher Education, Research and Innovation (for APAFIS permit numbers for mouse lines, see below).

In this study, we performed extensive phenotyping of the enteric nervous system in a genetically engineered mouse model that we previously generated (Creuzet et al. 2016). This model enables the conditional ablation of *Lkb1* in pre-migratory neural crest cells through the combination of *Lkb1* floxed alleles (*Lkb1^F^*^/F^), the *HtPA-Cre* transgene, and the R26R reporter *lacZ,* and is referred to as LHR throughout the study. We also obtained mice with a p53 heterozygous deletion (Jacks et al. 1994) from L. Le Cam (Institut de Recherche en Cancérologie de Montpellier, France). These mice were crossed with LRH animals to create the new LRHP line (APAFIS #8409-2016111515135080v11, between 2018-2023) and generate LRHP^+/-^ and LRHP^-/-^ offspring. Breeding of LRH and LRP genotypes was maintained at the hTAG unit (APAFIS #10040-2017031614558233.v4 for LRH line between 2018-2023, #15468-2018060816282691.v3 for LRHP for 2019-2024, and #48576-2024030615478465.v6 for LRHP from 2024). Breeders were transferred to the IAB animal facility to generate LRH and LRHP study cohorts (APAFIS #14637-2016120816101556v9 between 2018-2023 and #45574-2023051113301236v17 from 2024). After crossing with p53 null mice (mixed background C57Bl6; 129Sv/J; DBA; white coat color), LRHP animals were then backcrossed (2 generations) with C57BL/6 individuals to maintain Agouti coat color to enable us identifying mutants based on coat depigmentation. The generation and genotyping methods have been described previously (Creuzet et al. 2016).

For motor tests, animals were gently held by the base of their tail and lifted for a maximum of 10 seconds as previously performed (Radu et al. 2019). Female mice with vaginal plugs the day after mating were considered 0.5-day pseudo-pregnant, and embryos were harvested at the necessary developmental stage for the study. Embryos were harvested from pregnant females euthanized by cervical dislocation, and E15.5 embryos were euthanized by decapitation. Neonates were weighed and monitored daily from day 5 post-birth, when individual identification was possible. N-acetylcysteine (NAC, Sigma-Aldrich) was administered to pregnant and lactating females in the drinking water at a dose of 0.4g/kg/day. Fresh NAC solutions were prepared daily, with concentration adjusted based on the weight of the heaviest female and the volume of water consumed the previous day. AICAR (5-aminoimidazole-4-carboxamide riboside, 50mg/kg, resuspended in PBS) was administered daily via subcutaneous injection to pregnant females from E9.5 to birth as previously described (Rodrigues et al. 2014; Wu et al. 2012) (APAFIS #43209-2023042614438244v6, from 2023). Mendelian distributions were calculated based on the genotypes of individuals included in the histological and molecular analyses conducted throughout the study, and were used to determine the lethality coefficient. Genotype frequencies were then compared to expected values using Chi^2^ tests (Excel software, ref suppl meth).

### Cell culture conditions, siRNA transfection and antioxidant treatment

Mouse immortalized neural crest stem cells (JoMa1.3) (Maurer et al. 2007) were cultured as previously described for progenitors and glial derivatives (Radu et al. 2019). The differentiation of these cells into neural cells was induced by first coating the culture surfaces with Poly-L-Lysine (10 μg/mL, 1h at 37°C), followed by fibronectin. BMP2 (50 ng/mL; Peprotech, 120-02) was then added to the JoMa1.3 culture medium for a period of 6 days, with medium changes every two days. *Lkb1* knockdown was performed using siRNAs from Dharmacon (ON-TARGETplus Mouse *Stk11* SMART Pool and ON-TARGETplus Non-targeting Pool as control). siRNAs were transfected using jetPRIME (Ozyme) or RNAiMax lipofectamine (Invitrogen) with 60 nM siRNA final. For progenitor cells, experiments were performed 72 hours after transfection. For neural and glial differentiated JoMa1.3 cells, two cycles of 96 hours of *Lkb1* knockdown were performed. Biochemical dosage of alanine, MitoTracker staining and Seahorse analyses in neuronal NCCs were carried out as previously described (Radu et al. 2019). N-acetylcysteine (NAC, Sigma-Aldrich) antioxidant treatment was performed during glial specification by treating cells with 12 μM NAC the last 6 days of the protocol, and the medium was changed every 2 days. Normality of the data and homoscedasticity of the variances were checked, and statistical differences were evaluated using two-tailed t-test.

### Histology, Immunohistochemistry and immunofluorescence staining

Digestive tract samples were fixed in 4% formaldehyde (Sigma-Aldrich), dehydrated and embedded in paraffin. Sections for histology were stained with Harris Hematoxylin and Eosin (Sigma-Aldrich) according to the supplier’s protocol and as previously described (Creuzet et al. 2016).

For immunofluorescence, sections and cells cultivated on glass coverslips were treated with standard protocols using primary antibodies as described in Table 2. Secondary antibodies used were from Jackson ImmunoResearch (Donkey anti-Mouse Alexa-488, Donkey anti-Mouse Cya3, Donkey anti-Rabbit Alexa-488, Donkey anti-Rabbit Cya3, Donkey anti-Rabbit Cya2 and Goat anti-Chicken A488). For tissue sections, nuclei were stained with Hoechst 33342 (10 μg/ml, Invitrogen).

For JoMa1.3 cells, nuclei were stained with Hoechst. Sections and cells were mounted in Mowiol. Acquisitions and analyses were performed using an AxioImager Z1 microscope equipped with an AxioCam MRm camera, an AxioImager M2 microscope, equipped with a Hamamatsu Orca R2 camera, or for confocal microscopy a LSM710 microscope (AxioObserver Z1 multiparameter NLO – LIVE7 – Confocor3) (Carl Zeiss Microscopy GmbH). Statistical differences in quantifications conducted on histological sections were evaluated by first checking the normality of the samples and then employing a two-tailed t-test, with the selection of homoscedastic or heteroscedastic variances based on the results obtained.

### Embryo immunostaining and clearing

*Lkb1* wildtype or conditional knockout E10.5 embryos were fixed in 4% formaldehyde (Sigma-Aldrich) and stored in ethanol 70%. After washing in PBS, embryos were permeabilized in a bath of PBSGT (PBS 1x, 0.2% gelatin, 0.5% Triton 100X) on a rocker at 37°C for 4 days. Then, they were incubated with primary antibodies (Table 2) diluted in PBSGT + 0.1% saponin. After washing in PBSGT, embryos were incubated in the secondary antibody solution for 3 days at 37°C. Next, we used iDISCO+ clearing modified procedures (Ertürk et al. 2012; Renier et al. 2014). All incubation steps were performed on a rocking shaker at 15 rpm using 50 ml tubes protected from light. Mouse embryos (E10.5) were embedded in 1.5% Phytagel (Sigma-Aldrich) in water and dehydrated in graded series (50%, 70%, 90% and 100%) of methanol diluted in H_2_O_2_ during 1 hour each step. This was followed by overnight incubation in methanol 100% and 2 hours of delipidation in dichloromethane (DCM, Sigma-Aldrich). Samples were cleared overnight in dibenzylether (DBE, Sigma-Aldrich) and then stored in brown glass vials (Rotilabo, Roth) filled with ethyl-cinnamate (ECI, Sigma-Aldrich), in the dark at room temperature.

### Light Sheet Microscopy and 3D imaging of embryos

Optically*-*cleared and fluorescently-stained mouse E10.5 embryo*s* were imaged in ECI using a lightsheet fluorescence microscope (UltraMicroscope II) with a 2x objective (WD= 6 mm with the dipping cap) and ImspectorPro software (LaVision BioTec GmbH, Miltenyi Biotec B.V. & Co. KG). The voxel size for the acquisitions was: x=1.21; y=1.21; z=2 μm.

3D images reconstructions were generated using Imaris software (version 9.6, Bitplane, Oxford Instrument), and the segmentation of mouse embryo digestive tract and peripheral nerves was performed using the contour surface technique to extract a 3D object and create a new channel. This new channel was subsequently processed using FIJI software. Segmented SOX10 and p-p53 staining were then analyzed with the Analyze Particles plugin (size: 2-20; circularity 0.5-1.0) to measure the surface area stained by SOX10 or p-p53. Eventually, the percentage of surface area covered by both staining was calculated.

For a higher spatial resolution, a unique adaptive optics confocal microscope (Nikon A1R) equipped with the deformable mirror module (AOS-micro, AlpAO) for the correction of geometrical aberrations (“ConfoBright” system) with a 40x/1.15 water immersion objective with a long working distance (WD 610 μm) was used to image embryonic digestive tracts (E10.5). Indeed, the deep confocal imaging of the cleared 3D sample requires long distance objectives and immersion media of, generally, different refractive index, resulting in optical aberrations. The “ConfoBright” microscope iteratively corrects these geometric optical aberrations both in excitation and detection light path prior to imaging, and restores locally the high diffraction-limited spatial resolution and the best possible photon collection efficiency. The driving metrics based on molecular brightness or on fluorescence intensity were used to optimise the adaptive optics in the isoplanatic patch of ca. 100 μm at each imaged depth. A graphical user interface was written in MATLAB (The MathWorks Inc.) and used during image acquisition with NIS-Elements (NIKON, Europe B.V.).

To improve the embryo SOX10^+^ images*, w*e used a deconvolution software developed for lightsheet microscopy *(Becker et al. 2019)*.

### Whole-mount TUBB3 immunostaining of the enteric nervous system and confocal imaging

The enteric neural network was visualized according to the protocol previously described (Radu et al. 2019). Briefly, embryonic (E15.5) and postnatal intestines (P0) were fixed for 2 hours in 4% formaldehyde (Sigma-Aldrich) at 4°C and washed in PBS. Samples were surfixed and cleared 4h in methanol/DMSO (4:1). After washing twice in PBT (phosphate buffer saline-0.1% Triton X-100) during 5 min, tissues were incubated overnight with TUBB3 and HuC/D antibodies (Table 2), diluted in PBT containing 1% BSA and 0.15% Glycine (PBG). After extensive washings, tissues were incubated overnight with secondary antibodies of adequate species and conjugated either with A568 or A647 fluorophores (1/500; Jackson ImmunoResearch). After washes in PBG, tissues were counterstained with the nucleus dye Hoechst 33342 and incubated in glycerol 80%. Samples were mounted extemporaneously in fresh glycerol 80% and imaged using the “Confobright” system. 3D image acquisitions were performed with a PlanApo 20x/0.75 objective to quantify both HuC/D-positive nuclei and the TUBB3-postive axonal network. Nuclei were segmented using the StarDist deep-learning-based approach (https://github.com/stardist/stardist) included in the FIJI software (Schindelin et al. 2012). Maximum intensity projections (MIP) of the z-stacks from the axonal network images were first processed to enhance local contrast (CLAHE) and binarized, and the non-staining areas created by the TUBB3 mesh were then measured using the Analyze Particles plugin.

### Metabolic profiling by ex vivo nuclear magnetic resonance (NMR) of digestive tracts

Gut metabolic profiling using ^1^H HRMAS NMR was adapted from described previously (Radu et al. 2019).

<u>Sampling</u>: the third most distal segment of the intestines from newborn pups (P0) was quickly dissected, cleaned of mesenteric tissue, placed in a 50μl zirconium rotor within a 1.5ml Eppendorf tube, and immediately snap-frozen in liquid nitrogen. On the day of the acquisition, 25μl of D_2_O was added to the sample prior acquisition.

<u>Data acquisition</u>: all HRMAS NMR spectra were acquired on a Bruker Biospin NEO spectrometer (IRMaGe, CEA Grenoble, France) at 500 MHz. ^1^H spectra were acquired using a Carr-Purcell-Meiboom-Gill (CPMG) pulse sequence, with an echo time of 30ms. Samples were spun at 4KHz and temperature maintained at 4°C. The acquisition of one ^1^H NMR spectrum consisted of 250 scans lasting 14 min. Residual water signal was pre-saturated during the 1.7-s relaxation delay time.

<u>Data processing</u>: after Fourier transformation and manually phasing with the Bruker software Topspin version 4.0, further pre-processing steps (baseline correction, alignment, bucketing) were performed using NMR ProcFlow version 1.4 online (http://nmrprocflow.org). The spectra were then segmented in 0.001 ppm buckets between 0 and 8.5 ppm with exclusion of residual water peak. Each bucket was normalized to the sum of all buckets per spectrum. Metabolite relative concentrations were finally calculated as the sum of all buckets corresponding to a given peak with no spectral overlapping.

<u>Multivariate statistical analysis</u>: the buckets were loaded into the SIMCA v17 (Malmö, Sweden) software. We initially applied unsupervised Principal Components Analysis (PCA) to obtain an overall representation of sample distribution. Subsequently, we employed Orthogonal Partial Least Squares Discriminant Analysis (OPLS-DA) to identify metabolites that could discriminate between the two mouse strains (*Lkb1* WT or KO). The scores were plotted in 2D against the first two, components of the OPLS-DA models, while the loadings were plotted in 1D to simulate an NMR spectrum. In this representation, positive peaks corresponded to up-regulated metabolites in Lkb1 cKO compared to WT mice, while negative peaks indicated down-regulated metabolites. Within this “statistical spectrum”, each NMR variable was color-coded based on its correlation with group membership. Metabolites with a correlation ≥ 0.5 were deemed the most discriminative and subjected to additional univariate analysis. To do so, we summed the buckets within the spectral region corresponding to one peak of the metabolite, resulting in a relative amplitude measurement for the metabolite. Statistical difference was evaluated by first checking the normality of the samples, and then using two-way unpaired t-test.

### Western blot and Oxyblot

Analysis of proteins by Western blot from JoMa1.3 cells was performed as described previously (Radu et al. 2019), using antibodies indicated in Table 2. Fold change of p53 phosphorylation was calculated by the ratio of bands intensity of phosphorylated p53 over total p53 (normalized to actin signal), and siLkb1 condition was next indexed to the control condition. Statistical differences were determined by first examining the normality of the samples. If normality was achieved, a one-sample t-test was used. When normality was not met, a one-sample Wilcoxon test was employed.

Protein oxidation was assessed using an Oxyblot detection kit (Millipore-Sigma, S7150) and following the manufacturer’s instructions. Briefly, dissected digestive tracts from newborn pups were lysed in 50mM Tris-HCl, 1% NP40, 0.25% Sodium deoxycholate and proteases inhibitor cocktail. Protein concentration was measured using microBCA protein assay and concentration was adjusted to 3-4 μg/μl. SDS was added to obtained a 6% final concentration and derivatization of the samples was obtained by adding DNPH1x solution (v:v), incubating 5 minutes and adding neutralization buffer. Negative controls were obtained by preparing samples following the same protocol but without DNPH addition. 2-β-mercaptoethanol reducing agent was added at 0.4M before proceeding to electrophoresis and THBP detection using primary and secondary antibodies provided in the kit.

Oxyblot quantifications were performed using the Fiji plugin Band/Peak quantifications by normalizing oxidation signal (whole lane) over actin band intensity. Statistical analysis was performed using two-tailed t-test.

### ROS analysis by flow cytometry

ROS levels were assessed using the cellular ROS Detection Assay Kit from Abcam, according to the manufacturer’s instructions. In brief, JoMa1.3 progenitor or glial cells transfected with siRNA control or targeting Lkb1 were incubated with the DCF-DA probe (20 μM) for 30min at 37°C in serum-containing DMEM (10%) devoid of phenol red. DCF-DA positive cells were then analyzed using the Attune flow cytometer. Positive controls included cells treated with tert-butyl hydroperoxide (TBHP, 55 mM) or hydrogen peroxide (H_2_O_2_, 500 μM) for 1h (progenitors) or 1h (glial cells), while the negative control consisted of cells that were not exposed to DCF-DA. For N-acetyl-cysteine (NAC) treatment of glial cells, 12 μM NAC was added to glial cells at the beginning of culture.

### mRNA extraction, reverse transcription and quantitative polymerase chain reaction

Total RNA was extracted using TRIzol™ (Invitrogen™), and reverse-transcribed with the iScript™ Reverse Transcription Supermix (Bio-Rad). Quantitative polymerase chain reaction (qPCR) was performed using SsoAdvanced Universal SYBR Green Supermix (Bio-Rad) and CFX CONNECT (Bio-Rad). Sequences of primers used are given in Table 1. Housekeeping genes were glyceraldehyde-3-phosphate dehydrogenase (GAPDH) and actin beta (ACTB) for progenitors, and lipid droplet-regulating VLDL assembly factor (AUP1) and proto-oncogene NF-KB subunit (RELA) for glial cells; normalizations were performed using GAPDH for progenitors and AUP1 for glial cells. Gene expression from cKO samples was indexed to expression values of WT condition using the 2^−Δ ΔCt^ method. Statistical difference was tested using one-sample rank and Wilcoxon signed-rank tests.

**Table 1.**
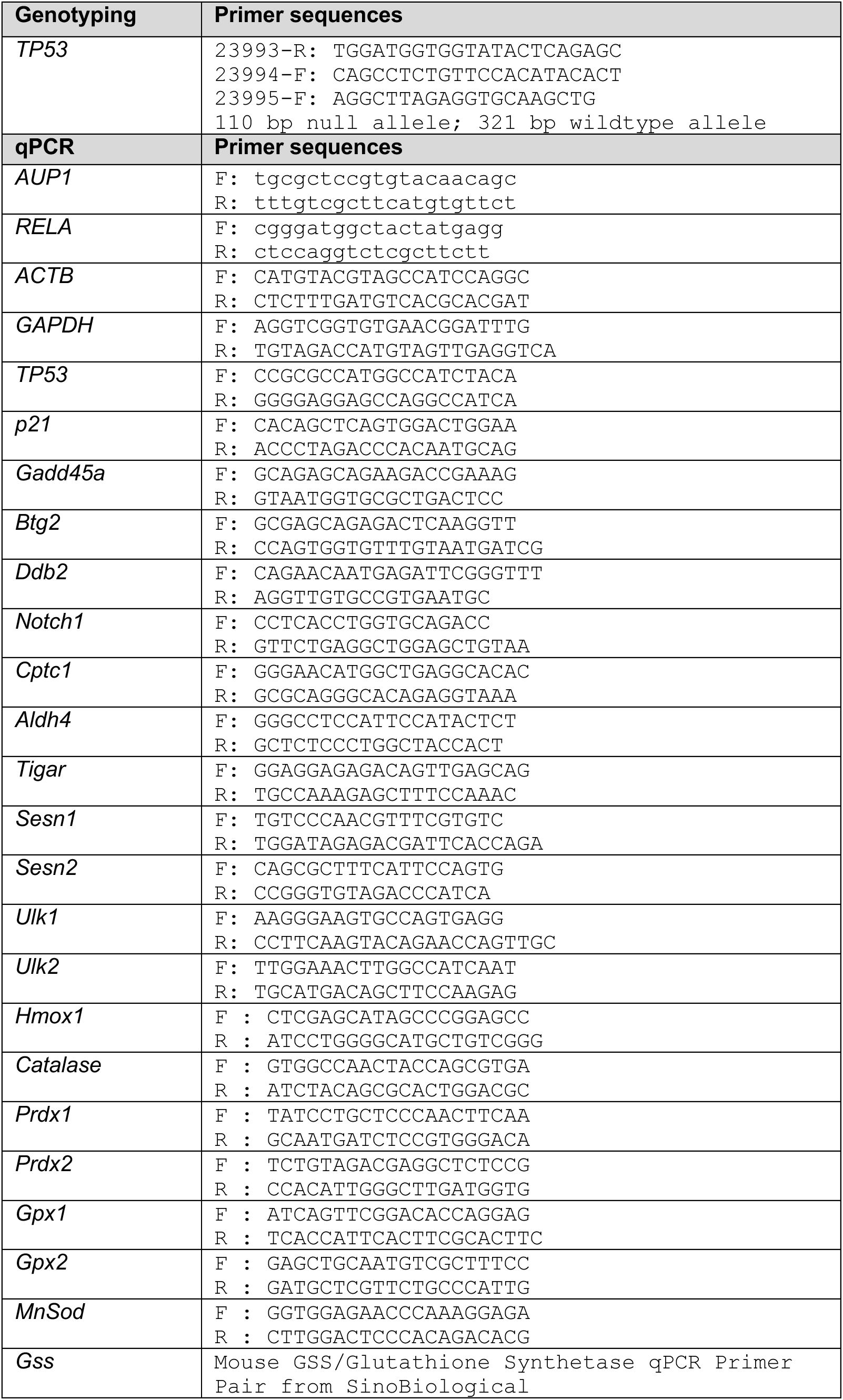
– Primers used for genotyping and RT-qPCR experiments. All primers target mouse genes. All primers are given from 5’ to 3’ left to right.

**Table 2.** – Antibodies used throughout the study.

| Technic | Primary antibody | Producing specie | Dilution | Embedding & Antigen retrieval | References |
| --- | --- | --- | --- | --- | --- |
| IHC | LKB1 D60C5 | Rabbit, polyclonal | 1/200 | Paraffin & Citrate | Cell signaling technology (#3031) |
| IF on tissue sections or whole mount staining | β-Galactosidase | Chicken, polyclonal | 1/500 | Paraffin & EDTA | Abcam (ab134435) |
|  | HUC/D | Mouse, monoclonal | 1/500 | Paraffin & NeoUltra | Molecular probe (A-21271) |
|  | HUC/D | Rabbit, polyclonal | 1/500 | Paraffin & NeoUltra | Abcam (ab184267) |
|  | GFAP | Rabbit, polyclonal | 1/200 | Paraffin & EDTA | Molecular probe (OPA1-06100) |
|  | GFAP | Chicken, polyclonal | 1/500 | Paraffin & EDTA | Abcam (ab4674) |
|  | TUBB3 (Tuj1) | Mouse, monoclonal | 1/1000 | Paraffin & NeoUltra | Covance (MMS-435P) |
|  | Phospho-S6RP (Ser235/236) | Rabbit, monoclonal | 1/1000 | Paraffin & NeoUltra | Cell signaling technology (#9661) |
|  | SOX10 A2 | Mouse, monoclonal | 1/1000 | Paraffin & NeoUltra | Santa Cruz (sc365691) |
|  | phospho-p53 (Ser15/18) | Rabbit, monoclonal | 1/200 | Paraffin & NeoUltra | Gene Tex (GTX 132995) |
|  | activated caspase 3 | Rabbit, polyclonal | 1/200 | Paraffin & Citrate | Cell signaling technology (#9661) |
|  | E-cadherin (clone 36) | Mouse, monoclonal | 1/200 | Paraffin & NeoUltra | BD transduction laboratories (#610181) |
| IF on cleared embryos or tissues | β-Galactosidase | Chicken, polyclonal | 1/500 | | Abcam (ab134435) |
|  | Peripherin | Rabbit polyclonal | 1/200 |  | Gene Tex (GTX 54569) |
|  | SOX10 A2 | Mouse, monoclonal | 1/1000 |  | Santa Cruz (sc365691) |
| IF on cultured cells | S100β | Mouse, monoclonal | 1/1000 | | Abcam (ab11178) |
|  | phospho-p53 (Ser15/18) | Rabbit, monoclonal | 1/1000 |  | Gene Tex (GTX 132995) |
| | γH2AX | Rabbit, monoclonal | 1/1000 | | Abcam (ab81299) |
| WB | phospho-p53 (Ser15/18) | Rabbit, monoclonal | 1/1000 |  | Gene Tex (GTX 132995) |
|  | Total p53 (CM5) | Rabbit, polyclonal | 1/1000 |  | Leica (NCL-p53-CM5p) |
|  | Lkb1 (D60C5) | Rabbit, polyclonal | 1/1000 |  | Cell signaling technology (#3047) |
|  | Actin | Mouse, monoclonal | 1/10000 |  | Millipore (MAB1501) |

### RNA sequencing

<u>RNA-Seq Library Construction and Sequencing</u>. RNA-Seq libraries were constructed with Optimal Dual-mode mRNA Library Prep Kit from Beijing Genomics Institute (BGI). A sequence read length of 150 bp was obtained by sequencing on the DNBSEQ T7 platform using the pair-end method (30M clean reads per library).

<u>Data Quality Control</u>. The raw data were filtered, low-quality sequence data removed, and quality controlled with SOAPnuke v2.3. The clean data were saved in FASTQ format for subsequent bioinformatics analysis.

<u>Genome and gene alignments</u>. The clean reads were aligned to the reference genome sequence *Mus musculus* (version: GCF_000001635.27_GRCm39) using HISAT2 software (v2.0.4). The software Bowtie2 v2.2.5 was used to align the clean reads to the reference gene sequence (transcriptome).

<u>Gene Expression Quantification</u>. RSEM v1.2.8 was used to calculate the gene expression levels in each sample. Fragments per kilobase of exon model per million mapped fragments (FPKM) values were used to estimate the level of gene expression. The genes with a mean FPKM > 0.5 were considered expressed in the group.

<u>Identification of Differentially Expressed Genes (DEGs)</u>. DEGs between the two groups were identified using DESeq2 (v1.4.5). Gene expression differences were calculated, and significant DEGs were defined as those meeting the thresholds of |log₂(FoldChange)| > 0.5 and Q-value < 0.05. DEG analysis was performed using the Dr.Tom Multi-Omics Data Mining System (BGI).

<u>GO and KEGG Pathway Enrichment Analyses.</u> The key biochemical and biological functions associated with the identified DEGs, along with the relevant metabolic and signal transduction pathways, were analyzed using the Gene Ontology database (http://geneontology.org/) and the KEGG database (http://www.kegg.jp/). Pathways enriched with at least two genes were considered for downstream analysis.

## Acknowledgments

The authors wish to thank Sylvie Dufour and Laurent Le Cam for sharing mouse lines and providing scientific advice for our project. We also thank Maud Sweitzer and Renaud Blervarque for their excellent technical assistance in *in vitro* analyses. We are deeply grateful to Arlette Lavigne from the IAB animal facility for her excellent care in housing and breeding the animals involved in this study, ensuring their well-being. We would also like to sincerely thank the UAR hTAG technical staff for dedicated care of the animals, ensuring the maintenance of the breeding colonies of the mouse strains used in this study. In particular, we are very grateful to Candice Hackney for her outstanding support in maintaining the mouse lines. Additionally, we acknowledge the MicroCell facility (GIS IBiSA, ISdV, IAB) for excellent support with various microscopy techniques throughout the study. The MicroCell facility is a member of the national infrastructure France-BioImaging.

## Author contributions

AL, ST, CT, and MB conceived the project, designed the studies, conducted most experiments, and analysed and interpreted the data. ST and MB developed the HtPA-Cre Lkb1 cKO mouse model. AL, JA, ST and CT conducted the animal experiments (breeding, genotyping and dissections). AL, ST, FA, JA and CT performed histological analyses for phenotypic characterization. BH performed all analyses of Mendelian distributions. AL, FA and CT carried out 3D lightsheet and confocal microscopy analyses, with FA and AG performing all 3D reconstructions, image analyses, and quantifications. JA and FF performed NMR acquisitions, quantifications and analyses with CT. AGR, MMA, ST, and CT conducted *in cellulo* experiments and analyses. JM provided the JoMa1.3 cell line. AL, FA, FF, ST, BF and CT wrote the manuscript, with all co-authors contributing to its final version.

## Figure legends

**Figure S1. Enteric nervous system formation and validation of the LRH model**.

Related to Fig.1 and 2.

**A.** Schematic representation of the digestive tract indicating the main areas of interest in this study: third most distal part of the intestine and the distal moiety of the colon used for transverse sections and histological analyses. **B.** Illustrations indicative of the migration paths followed by enteric progenitors at different embryonic days. At E10.5, enteric progenitors reach the midgut; at E11.5 they reach the cecum; grey arrow indicate both routes undertaken by enteric progenitors at this stage: rostro-caudal migration through the cecum and trans-mesenteric migration from the midgut to the hindgut. By E15.5, the whole digestive tract has been colonized. **C.** Timeline of birthdate of neurons and glial cells over the course of digestive tract colonization by enteric progenitors. **D,E.** Representative images of TUBB3– and HuC/D-positive immunostaining of enteric neurons (D) or GFAP-positive staining (E) in submucosal ganglia. Transverse sections of distal intestine of WT animals (P0). Ho: Hoechst. **F.** IF Representative images of HuCD immunofluorescence on proximal colon sections of WT and LRH P0 individuals. Quantification is shown and two-tailed t-test value is indicated.

**Figure S2.** Related to Fig.3

**A**. Representative TUBB3 labeling on whole mount digestive tracts of WT and LRH mice at E12.5. Arrows indicate the migration front of enteric progenitors. The LRH digestive tract is schematized (right) and the front migration is shown (arrow in the colon area) st.: stomach. **B**. Representative images of a WT embryo at E10.5 at dissection. After immunofluorescence labelling, embryos are embedded in Phytagel due to their small size (middle picture) and then cleared using 3DISCO protocol (bottom picture). **C**. 3D confocal images (ConfoBright) of peripherin and β-Gal labeling of E10.5 cleared embryos. The vagal area is zoomed on the right. Cranial nerves IX (glossopharyngeal), X (vagal), XI (accessory) and XII (hypoglossal) are indicated. β-Gal labeling is localized in the area of neural crest cell domains. White squares on the digestive tract indicate the zoomed area used to extract pictures for figure 3E. **D**. Confocal 3D images focusing on the head of Peripherin and β-Gal labeled E10.5 cleared embryos. β-Gal staining overlap with neural crest cells territories in the face.

**Figure S3. AICAR treatment did not prevent the defects in the enteric neuronal network caused by *Lkb1* loss.** Related to Fig.3

**A**. Schematic representation of the procedure used for AICAR subcutaneous administration to pregnant females and immunofluorescence analysis of the digestive tract of newborns. **B**. Maximum Intensity Projections (MIP) of adaptive optics confocal acquisitions (z-stack of 20 images z-step = 1.1 μm) of whole-mount digestive tracts of P0 mice stained for the axonal network (TUBB3, red) and neuronal nuclei (HuC/D, blue) in WT and LRH mutant mice (distal intestine). Controls consist of newborn from non-injected females (NT) or injected in PBS in which AICAR was resuspended. **C**. Quantification of the density of HuC/D-positive nuclei after segmentation, showing no rescue after AICAR treatment. WT-NT n=6, WT-PBS n=3, WT-AICAR n=6, LRH-NT n=3, LRH-AICAR n=6. **D.** Distribution of neuron nuclei density depending on their size measured by nuclei area. Densities shown on graph represent the normalized frequency of HuC/D-positive areas. LRH and LRH treated with AICAR showed increased small size nuclei compared to WT. Total nuclei counted (from n independent individuals): WT-NT=17396 (n=5), WT-AICAR=10709 (n=4), LRH-NT=3894 (n=3), and LRH-AICAR=9569 (n=6). **E.** Representative images on distal intestine illustrating MIP of adaptive optics confocal acquisitions (z-stack of 20 images over 22 μm total) to visualize the axonal enteric network using TUBB3 labelling (top) and non-innervated (TUBB3-negative) areas (bottom). **F.** Quantification of TUBB3-negative areas is shown. The marked increase of non-innervated areas upon *Lkb1* inactivation (LRH) was not rescued by AICAR treatment. WT-NT n=4, WT-PBS n=3, WT-AICAR =6, LRH-NT n=3, LRH-AICAR n=6. Statistical comparisons were performed using unpaired two-tailed t-tests (C, F). ns: non-significant.

**Figure S4.** Related to Fig.4

**A**. Representative pictures of E-cadherin (E-cad) immunolabeling on transverse sections of distal intestines showing that epithelial junctions are not altered in the LRH distal intestine at P0. Zoom on the intestinal epithelium is shown (right). WT and LRH n= 4. **B**. Metabolites detected in tissue using HRMAS NMR at 500 MHz. Assignemnt of peaks is provided in table in D. Representative ^1^H HRMAS NMR spectra obtained on digestive tracts of E13.5 (black) and P0 (blue) individuals. Assignment of resonances is identical in WT and LRH spectra. Abbreviations used for the attributions are detailed in table in D. **C**. S-line indicating metabolites that are upregulated (positive values) or downregulated (negative values) in LRH KO mice compared to WT controls at P0. 1D loading plot of the OPLS-DA built with P0 spectra, which looks like a 1D NMR spectrum. However, in this representation, the direction of peaks indicates whether metabolites are upregulated (positive) or downregulated (negative) in LRH digestive tracts compared to those in WT (P0 animals). The loading plots are color-codes based on their discriminative power. Assignments of some metabolites (from B) are reported into the loading plot to illustrate the most disturbed metabolites upon *Lkb1* loss. **D.** From left to right: Assignment of resonances to metabolites; Chemical shift (d, in ppm) of the peak used for quantification; sense of variation of KO vs WT at 13.5 and P0. List of identified metabolites into the ^1^H HRMAS NMR spectra, with identical (=), up-(+) and down-regulation (-) in LRH mice. Parts per million (ppm) values for each metabolite are indicated. These values correspond to the chemical shift of the peaks used for quantifications and univariate statistical analyses, rather than the chemical shift of the entire molecule. **E**. Relative amplitude of identified metabolites at E13.5 showing no statistical differences between WT animals and LRH mutants (t-test).

**Figure S5.** Related to Fig.5

**A**. Representative Western blots showing efficient *Lkb1* knockdown in neural crest progenitors and glial derivatives (JoMa1.3 cell line) upon *Lkb1* knockdown by siRNA (siLkb1) compared to scrambled siRNA (siCont.). **B.** Relative expression of several p53-target genes obtained on progenitor JoMa1.3 cells *in vitro* under Lkb1-targeting siRNA (siLkb1) and indexed to control condition (siCont/siLkb1 ratio). **C**. DNA damage labeling using antibody recognizing phosphorylated form of the Histone 2A variant (“H2AX) on glial JoMa1.3 cells *in vitro* under Lkb1-targeting siRNA (siLkb1) condition compared to control condition (siCont.). DNA damage increased upon *Lkb1* knockdown. n=3. **D**. Schematic representation of NAC treatment in drinking water of pregnant and lactating females and follow up of newborn pups. **E**. Weight gain curve of WT and LRH newborn pups is shown after NAC. WT n=13, WT/NAC n=14; LRH n=3, LRH/NAC n=2. Statistical analysis was conducted using mixed-effects models (a similar approach to a two-way ANOVA, except for datasets that are not equivalent between groups), followed by Tukey’s multiple comparison test, with p-values reported. **F,H**. Read count for Lkb1 (F) and p53 (H) gene expression measured in bulk RNAseq analyses in progenitor cells. **G**. Table of gene set enrichments of various cellular processes from differentially expressed genes from the RNAseq analysis in control of Lkb1-knockdown progenitors.

**Figure S6.** Related to Fig.6

**A**. Schematic representation of neural differentiation from JoMa1.3 progenitors. **B**. Experimental timeline of the culture protocol to derive neural cell types. **C**. Immunofluorescence images showing high neuronal (TUBB3) but no glial (GFAP) staining to validate the differentiation of neural derivatives in culture. **D**. Representative western blot analysis of neuronal cells validating *Lkb1* knockdown by siRNA (siLkb1) compared to scrambled siRNA (siCont.). **E.** Phase contrast images illustrating morphological differences between progenitors and their neuronal derivatives, as well as changes induced by Lkb1 knockdown (siLkb1). **F**. Immunofluorescence using TUBB3 neuronal marker identified lack of neuronal differentiation following Lkb1 knockdown. **G**. Western blot analyses of Lkb1 downstream signaling with AMPK (middle) and mTOR activity (right) quantifications. n=7 (AMPK); n=6 (S6RP). **H.** Intracellular alanine measurements in neural cells did not show any accumulation upon Lkb1 knockdown compared to progenitors and glial cells as previously published. **I.** Lkb1 loss triggered mitochondria mass increase in neurons as quantified by cell sorting analyses using live MitoGreen staining. **J**. Mitochondrial topology was impaired upon *Lkb1* knockdown with more tubular network compared to neuronal control cells. **K**. Seahorse analyses of oxygen consumption (OCR) showed no statistical differences between control and Lkb1-knockdown neuronal cells. **L**. Western blot analyses of phosphorylated p53 on Serine 18 (p-p53^Ser18^) following *Lkb1* knockdown in NCC-derived neurons show no detectable increase (graph on the right). n=5.

**Figure S7** – Related to Figure 7

**A**. Epifluorescence microscopy showing phosphorylated p53 labeling on transverse intestinal sections of E15.5 mice. Quantifications of spots of high phosphorylation signal (left, significant difference) and global phosphorylation intensity (right, not significant difference) are shown. WT and LRH n=4. **B**. Coimmunofluorescence of phosphor-p53 with glial (GFAP) and neuronal (TUBB3) markers show hyperphosphorylated p53 in both enteric neurons and glial cells. **C**. Same as in A on distal intestine of P0 animals. Density of highly intense phosphorylated p53 spots in ganglia is quantified (graph) showing increased number in LRH mutants. WT: n=3; LRH: n=4. **D**. TUBB3 and active caspase 3 (Casp3) co-immunofluorescence on whole mount digestive tracts of E13.5 mouse embryos. Active caspase 3 events were increased in LRH mice. **E**. Epifluorescence microscopy showing active caspase 3 labeling (Casp3) on transverse sections of distal intestine from P0 mice. No events were detected. WT and LRH n=4.

**Figure S8** – Related to Fig.7

**A**. List of all genotypes generated by crossing LRH mice with *p53* knockout mice, including the number and color codes. The diagram outlines the main breeding strategy used to produce LRH, LRHP^+/-^ and LRHP^-/-^ mice. The expected frequencies for key genotypes are indicated. A total of 703 offspring were generated across all listed genotypes for the various analyses conducted throughout the study, with 82 animals exhibiting with a detrimental phenotype. **B**. Weight curve of pups from 5 to 16 days of age with *Lkb1* conditional ablation (LRH), with or without ablation of one *p53* allele (LRHP^+/-^), compared to wildtype (WT) controls. The number of pups at birth was: WT n=17; LRH n=3; LRHP^+/-^ n=6. Statistical comparisons were performed using two-way ANOVA followed by Tukey’s multiple comparison test, with p-values reported for significant differences. No significant differences were found between the LRH and LRHP groups. **C.** Loss of hind limb extension reflex in 10-day-old WT and LRHP+/-pup as evidenced by hind limb clenching when lifted by the tail (motor test). **D.** Populations at all embryonic stages followed a Mendelian distribution: *Lkb1^F/F^ mice or Lkb1^+/F^,* male or female, were crossed with *Lkb1^+/F^ HtPA-Cre^+/tg^* mice of the opposite sex. Data from litters where the genotype of one parent was not confirmed were discarded. The number of *Lkb1^F/F^*embryos whether *HtPA-Cre^+/tg^* or ^+*/+*^ (respectively noted LRH and WT), was compared to the expected numbers according to Mendelian’s law, using Chi^2^ tests. Age is in days *post coitum*. ns: non-significant. **E.** To extend this analysis to P0 animals, all litters with confirmed parental genotypes were included, totaling 703 offspring. The table summarizes the null hypotheses tested (“questions asked”), the genotypes analyzed (as shown in panel A), the number of pups evaluated, Chi^2^ test p-values, and whether each null hypothesis was validated (YES) or rejected (NO). **F**. Focus on a specific analysis from Table E. The lethality coefficient for homozygous *Lkb1* loss was calculated using the following formula: *theorical number = Mendelian expected number *(100-lethality)/100*. This coefficient was individually optimized to the closest integer values that best fit the experimental data for genotypes 7 and 13, corresponding to the effects of conditional ablation of one or two *Lkb1* alleles, respectively. The results of the Chi^2^ test applied and the associated p-value are shown for Mendelian inheritance (left) and after accounting for the percentage of lethality (right).

**Figure S9** – Related to Figure 7

Maximum Intensity Projections (MIP) of adaptive optics confocal acquisitions (z-stack of 20 images z-step = 1.1 μm) of whole-mount digestive tract of P0 mice to visualize neuronal nuclei (HuC/D), nuclei segmentation, the axonal enteric network (TUBB3), and the analysis of non-innervated areas. Representative images in distal intestine illustrating the marked reduction of the axonal network upon *Lkb1* inactivation (LRH) but no rescue upon ablation of *p53* in LRH mutants.

## Supplemental Methods

### Statistical analysis of LHR and LHRP survival at birth

To analyze the survival of P0 LHRP animals, litters depicted in Figure S6A with confirmed parent genotypes, totaling 703 offspring, were included in a global statistical analysis. The hypothesis of Mendelian inheritance for this population was rejected (FigureS6E, line 1). We then analyzed the distribution of WT and intermediate genotypes (i.e. without a complete knockout of either p53 or Lkb1), excluding LKB1^+/+^ genotypes due to their low frequencies (FigureS6E, line 2). Mendelian inheritance was not ruled out, and we concluded that no strong genetic linkage interfered with the genotype distribution in these 534 offspring, which was extrapolated to all 703 pups in the same litters (note that the chromosomal location of the HtPA-Cre allele is unknown).

Next, we assessed the effect of p53 knockout in the absence of Lkb1 floxed alleles, focusing on 441 offspring. The Chi^2^ test was not significant (FigureS6E, line 3), indicating that p53 inheritance did not deviate significantly from the expected Mendelian distribution.

Similarly, the effect of Lbk1 inactivation driven by HtPA-mediated Cre expression was analyzed, excluding offspring with one or two p53 null alleles. A significant (p<10^-6^) excess of pups with an intact Lkb1 allele was observed in the 342 pups analyzed (Figure S6E, line 4). Details of the expected numbers and estimated lethality are provided in Figure S6E.

When comparing the number of LKB1^+/F^ Cre^+/tg^ to LKB1^+/F^ Cre^+/+^ pups, the Chi^2^ test did not rule out Mendelian inheritance (Figure S6F). However, for Lkb1^F/F^ Cre^+/tg^ pups, the Chi^2^ was significant, indicating that the number of LRH homozygous individuals obtained was significantly different from the expected number. The lethality coefficient was calculated using the following formula: theorical number = Mendelian expected number *(100-lethality)/100.

The lethality coefficient for LRH P0 animals was found to be 58%, and the theorical numbers within the LKB1 subpopulation were recalculated using this coefficient, optimized to the nearest integer, and no statistical difference was found (Figure S6F). Next, we assessed whether p53 loss modified Lkb1 inheritance in 62 pups but found no statistical difference (Figure S6E, line 5), indicating that p53 loss neither rescued nor worsened Lkb1-driven survival. Finally, recalculated distributions of LRH animals within the global population (703 pups), accounting for the 58% of LRH pups, also showed no statistical difference (Figure S6E, line 6), confirming that p53 loss did not affect the inheritance frequency of the Lkb1 mutant allele.

## Bibliography

1. Anderson RB, Newgreen DF, Young HM. 2006. Neural crest and the development of the enteric nervous system. Adv Exp Med Biol 589: 181–196.

2. Baas AF, Kuipers J, van der Wel NN, Batlle E, Koerten HK, Peters PJ, Clevers HC. 2004. Complete polarization of single intestinal epithelial cells upon activation of LKB1 by STRAD. Cell 116: 457–466.

3. Barriga EH, Franze K, Charras G, Mayor R. 2018. Tissue stiffening coordinates morphogenesis by triggering collective cell migration in vivo. Nature. https://www.nature.com/articles/nature25742 (Accessed February 20, 2018).

4. Becker K, Saghafi S, Pende M, Sabdyusheva-Litschauer I, Hahn CM, Foroughipour M, Jährling N, Dodt H-U. 2019. Deconvolution of light sheet microscopy recordings. Sci Rep 9: 17625.

5. Beirowski B. 2018. The LKB1-AMPK and mTORC1 Metabolic Signaling Networks in Schwann Cells Control Axon Integrity and Myelination: Assembling and upholding nerves by metabolic signaling in Schwann cells. BioEssays 1800075.

6. Beirowski B, Babetto E, Golden JP, Chen Y-J, Yang K, Gross RW, Patti GJ, Milbrandt J. 2014a. Metabolic regulator LKB1 is crucial for Schwann cell-mediated axon maintenance. Nat Neurosci 17: 1351–1361.

7. Beirowski B, Babetto E, Golden JP, Chen Y-J, Yang K, Gross RW, Patti GJ, Milbrandt J. 2014b. Metabolic regulator LKB1 plays a crucial role in Schwann cell-mediated axon maintenance. Nat Neurosci 17: 1351–1361.

8. Bhattacharya D, Azambuja AP, Simoes-Costa M. 2020. Metabolic Reprogramming Promotes Neural Crest Migration via Yap/Tead Signaling. Developmental Cell 53: 199–211.e6.

9. Bondurand N, Sham MH. 2013. The role of SOX10 during enteric nervous system development. Developmental Biology 382: 330–343.

10. Bourouh M, Marignani PA. 2022. The Tumor Suppressor Kinase LKB1: Metabolic Nexus. Front Cell Dev Biol 10: 881297.

11. Bowen ME, Attardi LD. 2019. The role of p53 in developmental syndromes. J Mol Cell Biol 11: 200–211.

12. Bowen ME, McClendon J, Long HK, Sorayya A, Van Nostrand JL, Wysocka J, Attardi LD. 2019. The Spatiotemporal Pattern and Intensity of p53 Activation Dictates Phenotypic Diversity in p53-Driven Developmental Syndromes. Dev Cell 50: 212–228.e6.

13. Charrier B, Pilon N. 2017. Toward a better understanding of enteric gliogenesis. Neurogenesis 4: e1293958.

14. Creuzet SE, Viallet JP, Ghawitian M, Torch S, Thélu J, Alrajeh M, Radu AG, Bouvard D, Costagliola F, Borgne ML, et al. 2016. LKB1 signaling in cephalic neural crest cells is essential for vertebrate head development. Developmental Biology 418: 283–296.

15. Erickson AG, Kameneva P, Adameyko I. 2023. The transcriptional portraits of the neural crest at the individual cell level. Semin Cell Dev Biol 138: 68–80.

16. Ertürk A, Becker K, Jährling N, Mauch CP, Hojer CD, Egen JG, Hellal F, Bradke F, Sheng M, Dodt H-U. 2012. Three-dimensional imaging of solvent-cleared organs using 3DISCO. Nat Protoc 7: 1983–1995.

17. Espinosa-Medina I, Jevans B, Boismoreau F, Chettouh Z, Enomoto H, Müller T, Birchmeier C, Burns AJ, Brunet J-F. 2017. Dual origin of enteric neurons in vagal Schwann cell precursors and the sympathetic neural crest. PNAS 114: 11980–11985.

18. Fischhuber K, Matzinger M, Heiss EH. 2020. AMPK Enhances Transcription of Selected Nrf2 Target Genes via Negative Regulation of Bach1. Front Cell Dev Biol 8: 628.

19. Gupta R, Liu AY, Glazer PM, Wajapeyee N. 2015. LKB1 preserves genome integrity by stimulating BRCA1 expression. Nucleic Acids Res 43: 259–271.

20. Holland AM, Bon-Frauches AC, Keszthelyi D, Melotte V, Boesmans W. 2021. The enteric nervous system in gastrointestinal disease etiology. Cell Mol Life Sci 78: 4713–4733.

21. Jacks T, Remington L, Williams BO, Schmitt EM, Halachmi S, Bronson RT, Weinberg RA. 1994. Tumor spectrum analysis in p53-mutant mice. Curr Biol 4: 1–7.

22. Jessen KR, Mirsky R. 2019. Schwann Cell Precursors; Multipotent Glial Cells in Embryonic Nerves. Front Mol Neurosci 12: 69.

23. Kameneva P, Kastriti ME, Adameyko I. 2021. Neuronal lineages derived from the nerve-associated Schwann cell precursors. Cell Mol Life Sci 78: 513–529.

24. Kang Y-N, Fung C, Vanden Berghe P. 2021. Gut innervation and enteric nervous system development: a spatial, temporal and molecular tour de force. Development 148: dev182543.

25. Kastriti ME, Faure L, Von Ahsen D, Bouderlique TG, Boström J, Solovieva T, Jackson C, Bronner M, Meijer D, Hadjab S, et al. 2022. Schwann cell precursors represent a neural crest-like state with biased multipotency. EMBO J 41: e108780.

26. Kullmann L, Krahn MP. 2018. Controlling the master-upstream regulation of the tumor suppressor LKB1. Oncogene 37: 3045–3057.

27. Lasrado R, Boesmans W, Kleinjung J, Pin C, Bell D, Bhaw L, McCallum S, Zong H, Luo L, Clevers H, et al. 2017. Lineage-dependent spatial and functional organization of the mammalian enteric nervous system. Science 356: 722–726.

28. Lefèvre MA, Godefroid Z, Soret R, Pilon N. 2024. Enteric glial cell diversification is influenced by spatiotemporal factors and source of neural progenitors in mice. Front Neurosci 18: 1392703.

29. Lyengar P, Gandhi AY, Granados J, Guo T, Gupta A, Yu J, Llano EM, Zhang F, Gao A, Kandathil A, et al. 2023. Tumor loss-of-function mutations in STK11/LKB1 induce cachexia. JCI Insight 8: e165419.

30. Margolis KG, Cryan JF, Mayer EA. 2021. The Microbiota-Gut-Brain Axis: From Motility to Mood. Gastroenterology 160: 1486–1501.

31. Matzinger M, Fischhuber K, Pölöske D, Mechtler K, Heiss EH. 2020. AMPK leads to phosphorylation of the transcription factor Nrf2, tuning transactivation of selected target genes. Redox Biol 29: 101393.

32. Maurer J, Fuchs S, Jäger R, Kurz B, Sommer L, Schorle H. 2007. Establishment and controlled differentiation of neural crest stem cell lines using conditional transgenesis. Differentiation 75: 580–591.

33. Meek DW, Anderson CW. 2009. Posttranslational Modification of p53: Cooperative Integrators of Function. Cold Spring Harbor Perspectives in Biology 1: a000950– a000950.

34. Nekooie Marnany N, Fodil R, Féréol S, Dady A, Depp M, Relaix F, Motterlini R, Foresti R, Duband J-L, Dufour S. 2023. Glucose oxidation drives trunk neural crest cell development and fate. Journal of Cell Science 136: jcs260607.

35. Newbern JM. 2015. Molecular Control of the Neural Crest and Peripheral Nervous System Development. In Current Topics in Developmental Biology, Vol. 111 of, pp. 201–231, Elsevier https://linkinghub.elsevier.com/retrieve/pii/S0070215314000088 (Accessed September 11, 2023).

36. Niesler B, Kuerten S, Demir IE, Schäfer K-H. 2021. Disorders of the enteric nervous system — a holistic view. Nat Rev Gastroenterol Hepatol 18: 393–410.

37. Ollila S, Mäkelä TP. 2011. The tumor suppressor kinase LKB1: lessons from mouse models. J Mol Cell Biol 3: 330–340.

38. Pawolski V, Schmidt MHH. 2020. Neuron-Glia Interaction in the Developing and Adult Enteric Nervous System. Cells 10: 47.

39. Perera SN, Kerosuo L. 2021. On the road again: Establishment and maintenance of stemness in the neural crest from embryo to adulthood. Stem Cells 39: 7–25.

40. Pham TD, Gershon MD, Rothman TP. 1991. Time of origin of neurons in the murine enteric nervous system: sequence in relation to phenotype. J Comp Neurol 314: 789–798.

41. Pietri T, Eder O, Blanche M, Thiery JP, Dufour S. 2003. The human tissue plasminogen activator-Cre mouse: a new tool for targeting specifically neural crest cells and their derivatives in vivo. Dev Biol 259: 176–187.

42. Pooya S, Liu X, Kumar VBS, Anderson J, Imai F, Zhang W, Ciraolo G, Ratner N, Setchell KDR, Yoshida Y, et al. 2014. The tumour suppressor LKB1 regulates myelination through mitochondrial metabolism. Nat Commun 5: 4993.

43. Radu AG, Torch S, Fauvelle F, Pernet-Gallay K, Lucas A, Blervaque R, Delmas V, Schlattner U, Lafanechère L, Hainaut P, et al. 2019. LKB1 specifies neural crest cell fates through pyruvate-alanine cycling. Sci Adv 5: eaau5106.

44. Rahman AA, Ohkura T, Bhave S, Pan W, Ohishi K, Ott L, Han C, Leavitt A, Stavely R, Burns AJ, et al. 2024. Enteric neural stem cell transplant restores gut motility in mice with Hirschsprung disease. JCI Insight 9: e179755.

45. Raj S, Jaiswal SK, DePamphilis ML. 2022. Cell Death and the p53 Enigma During Mammalian Embryonic Development. Stem Cells 40: 227–238.

46. Rao M, Gershon MD. 2018. Enteric nervous system development: what could possibly go wrong? Nat Rev Neurosci 19: 552–565.

47. Renier N, Wu Z, Simon DJ, Yang J, Ariel P, Tessier-Lavigne M. 2014. iDISCO: A Simple, Rapid Method to Immunolabel Large Tissue Samples for Volume Imaging. Cell 159: 896–910.

48. Rodrigues SC, Pantaleão LC, Nogueira TC, Gomes PR, Albuquerque GG, Nachbar RT, Torres-Leal FL, Caperuto LC, Lellis-Santos C, Anhê GF, et al. 2014. Selective regulation of hepatic lipid metabolism by the AMP-activated protein kinase pathway in late-pregnant rats. *American Journal of Physiology-Regulatory*, Integrative and Comparative Physiology 307: R1146–R1156.

49. Ryter SW. 2022. Heme Oxygenase-1: An Anti-Inflammatory Effector in Cardiovascular, Lung, and Related Metabolic Disorders. Antioxidants 11: 555.

50. Schindelin J, Arganda-Carreras I, Frise E, Kaynig V, Longair M, Pietzsch T, Preibisch S, Rueden C, Saalfeld S, Schmid B, et al. 2012. Fiji: an open-source platform for biological-image analysis. Nat Methods 9: 676–682.

51. Seguella L, Gulbransen BD. 2021. Enteric glial biology, intercellular signalling and roles in gastrointestinal disease. Nat Rev Gastroenterol Hepatol 18: 571–587.

52. Shadfar S, Parakh S, Jamali MS, Atkin JD. 2023. Redox dysregulation as a driver for DNA damage and its relationship to neurodegenerative diseases. Transl Neurodegener 12: 18.

53. Sharkey KA, Mawe GM. 2023. The enteric nervous system. Physiol Rev 103: 1487–1564.

54. Shen Y-AA, Chen Y, Dao DQ, Mayoral SR, Wu L, Meijer D, Ullian EM, Chan JR, Lu QR. 2014. Phosphorylation of LKB1/Par-4 establishes Schwann cell polarity to initiate and control myelin extent. Nat Commun 5: 4991.

55. Sies H, Jones DP. 2020. Reactive oxygen species (ROS) as pleiotropic physiological signalling agents. Nat Rev Mol Cell Biol 21: 363–383.

56. Suzuki YJ, Carini M, Butterfield DA. 2010. Protein carbonylation. Antioxid Redox Signal 12: 323–325.

57. Tan I, Xu S, Huo J, Huang Y, Lim H-H, Lam K-P. 2023. Identification of a novel mitochondria-localized LKB1 variant required for the regulation of the oxidative stress response. J Biol Chem 299: 104906.

58. Thibert C, Lucas A, Billaud M, Torch S, Mével-Aliset M, Allard J. 2023. Functions of LKB1 in neural crest development: the story unfolds. Developmental Dynamics dvdy.581.

59. Ulisse V, Dey S, Rothbard DE, Zeevi E, Gokhman I, Dadosh T, Minis A, Yaron A. 2020. Regulation of axonal morphogenesis by the mitochondrial protein Efhd1. Life Sci Alliance 3: e202000753.

60. Wright CM, Schneider S, Smith-Edwards KM, Mafra F, Leembruggen AJL, Gonzalez MV, Kothakapa DR, Anderson JB, Maguire BA, Gao T, et al. 2021. scRNA-Seq Reveals New Enteric Nervous System Roles for GDNF, NRTN, and TBX3. Cellular and Molecular Gastroenterology and Hepatology 11: 1548–1592.e1.

61. Wu Y, Viana M, Thirumangalathu S, Loeken MR. 2012. AMP-activated protein kinase mediates effects of oxidative stress on embryo gene expression in a mouse model of diabetic embryopathy. Diabetologia 55: 245–254.

62. Xu H-G, Zhai Y-X, Chen J, Lu Y, Wang J-W, Quan C-S, Zhao R-X, Xiao X, He Q, Werle KD, et al. 2015. LKB1 reduces ROS-mediated cell damage via activation of p38. Oncogene 34: 3848–3859.

63. Zhou B, Feng C, Sun S, Chen X, Zhuansun D, Wang D, Yu X, Meng X, Xiao J, Wu L, et al. 2024. Identification of signaling pathways that specify a subset of migrating enteric neural crest cells at the wavefront in mouse embryos. Developmental Cell 59: 1689–1706.e8.

64. Zurkirchen L, Sommer L. 2017. Quo vadis: tracing the fate of neural crest cells. Curr Opin Neurobiol 47: 16–23.

