## supplemental figures for "The metabolic kinase LKB1 shapes the enteric nervous system by mitigating the oxidative stress and p53 activity"

Sup Movies :

- Movie1: ganglion confocal (associated with fig1)
- Movie2: zoom confobright SNE (z-stack) of E10.5 embryos
- Movie3: Sox10-positive area (z-stack) in digestive tract of E10.5 embryos
- Movie4: p-p53-positive area (z-stack) in digestive tract of E10.5 embryos

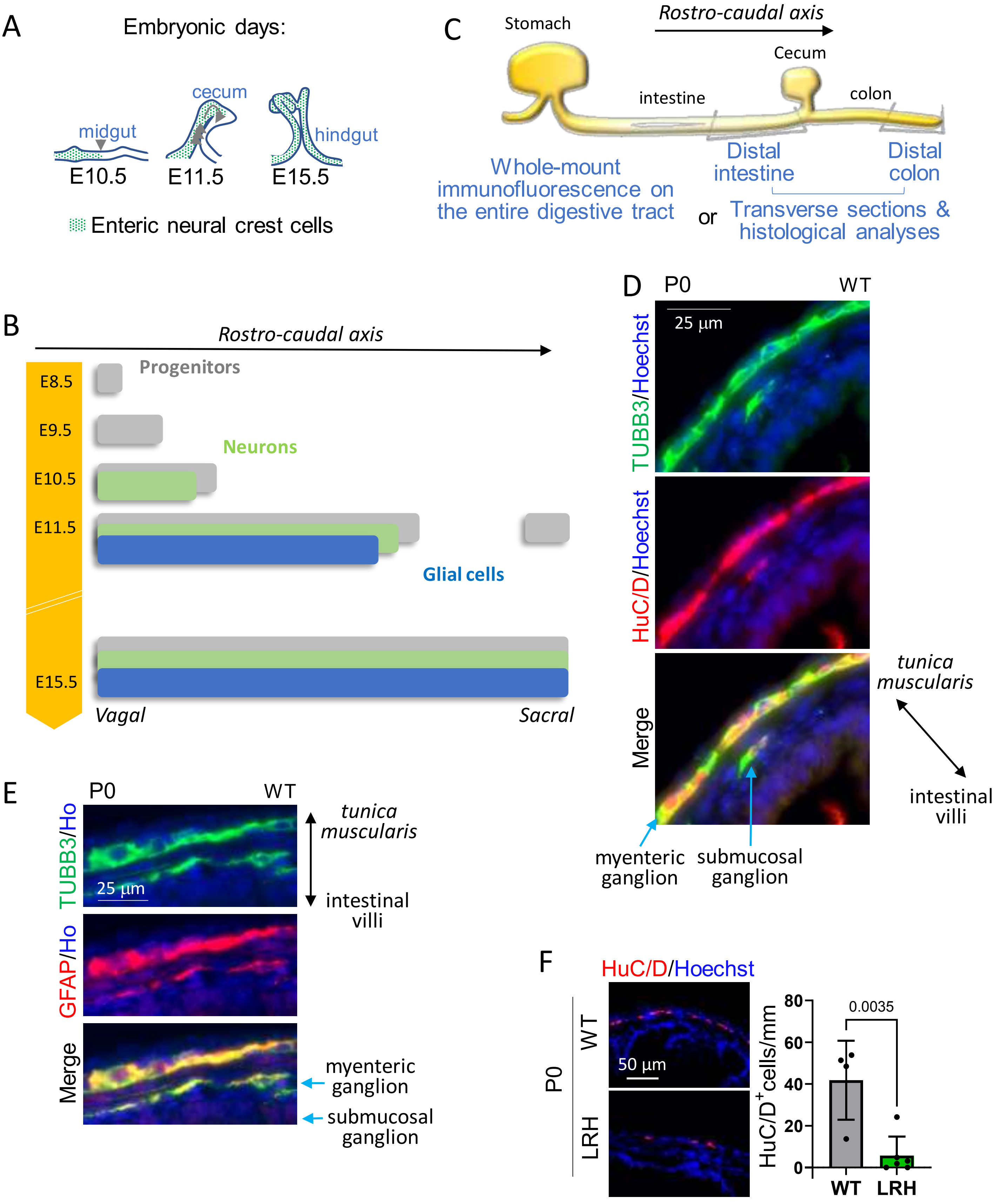


Fig. S1 Related to Fig. 1&2

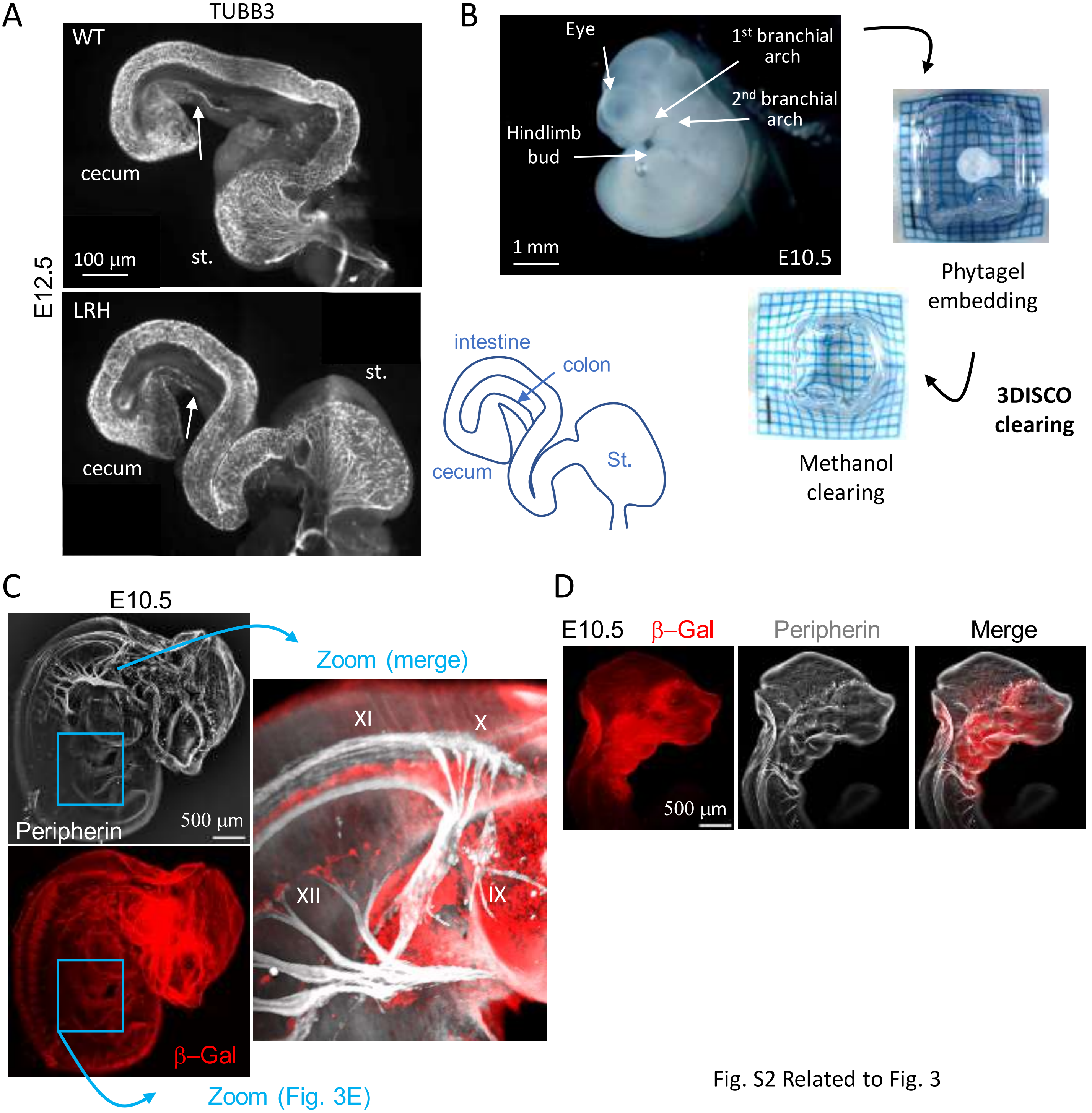


Fig. S2 Related to Fig. 3

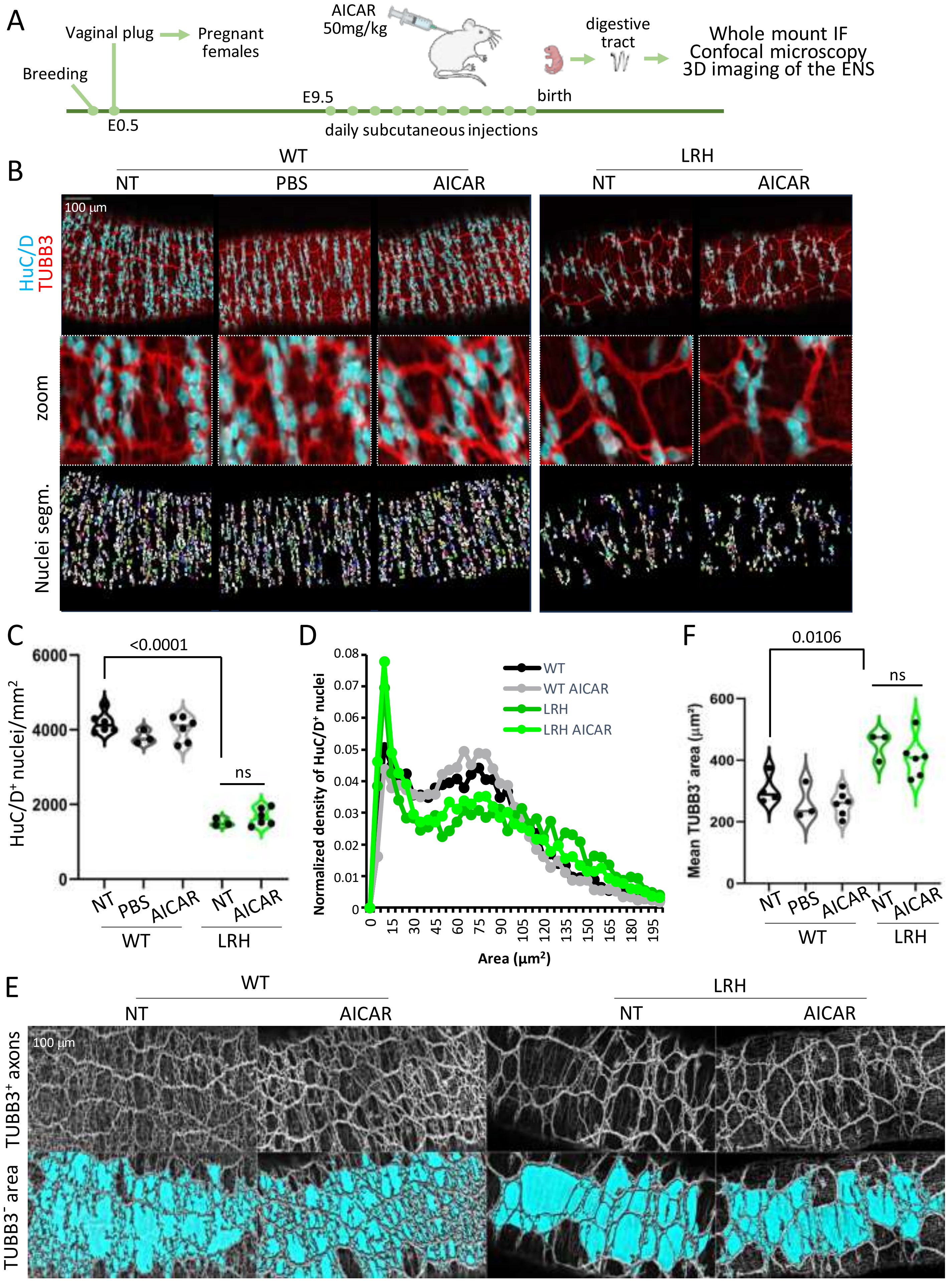


FigS3 related to Fig3. AICAR treatment during embryogenesis does not rescue the enteric neuronal phenotype

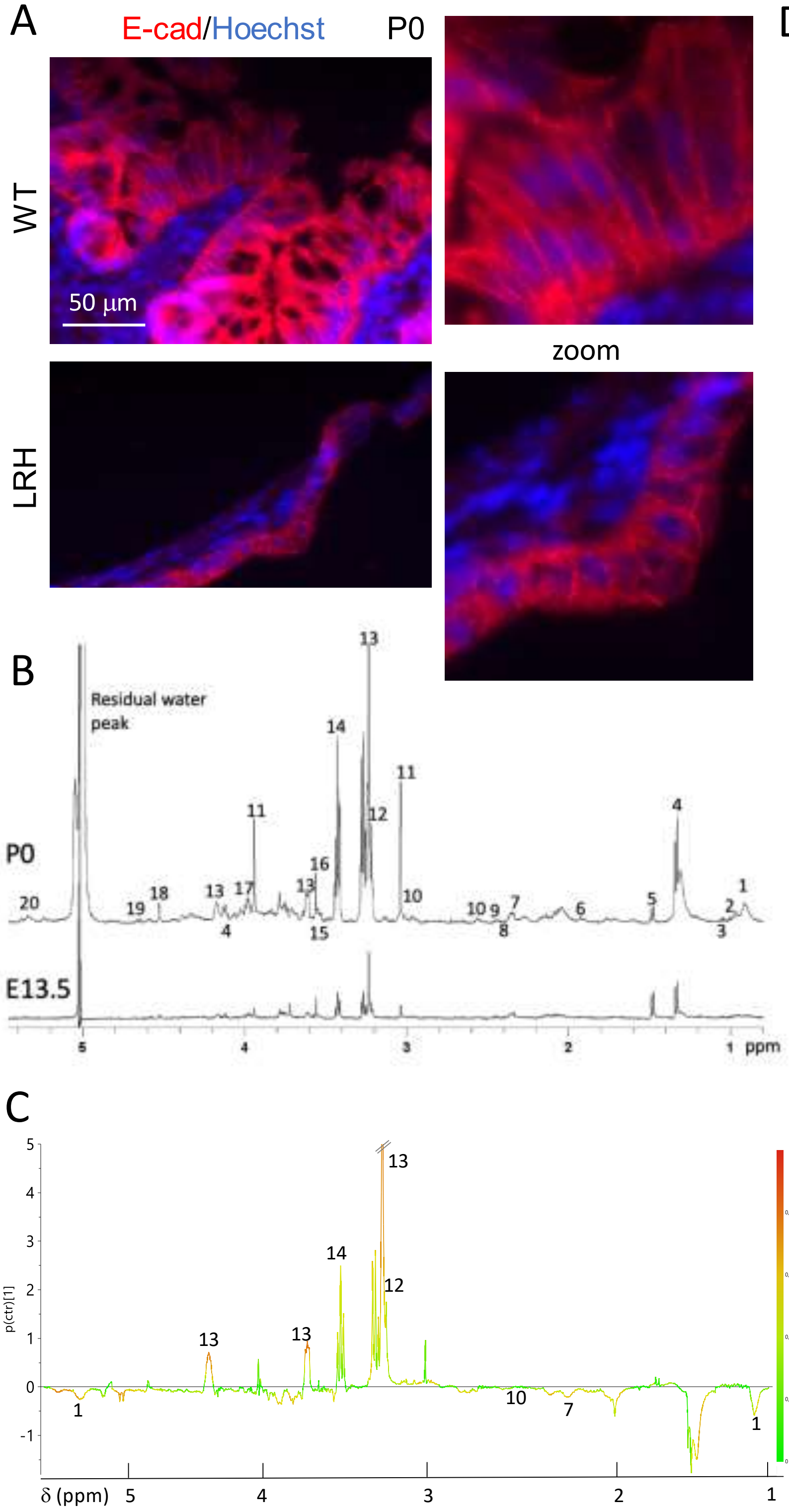


D

| N° | Metabolite | δ ppm | E13.5 | P0 |
|---|---|---|---|---|
| 1 | Lipids CH3 | 1.30 | | ↘ |
| 2 | Isoleucine | 1.02 | | = |
| 3 | Valine | 1,05 | = | = |
| 4 | Lactate | 4.123 | = | = |
| 5 | Alanine | 1.475 | = | = |
| 6 | Acetate | 1.93 | | = |
| 7 | Glutamate | 2.35 | = | ↘ |
| 8 | Succinate | 2.41 | | ↘ |
| 9 | Glutamine | 2.44 | = | = |
| 10 | Reduced Glutathione | 2.94 | | ↘ |
| 11 | Creatine/ Phospho-creatine | 3.03 | = | = |
| 12 | Choline | 4.16 | | ↗ |
| 13 | Phospho-choline | 3.23 | = | ↗ |
| 14 | Taurine | 3.42 | = | ↗ |
| 15 | Myoinositol | 3.536 | = | = |
| 16 | Glycine | 3.56 | = | = |
| 17 | Phospho-ethanol-amine | 3.98 | = | = |
| 18 | Ascorbate | 3.526 | = | = |
| 19 | α-Glucose | 5.266 | | = |
| 20 | Lipids-CH | 5.33 | | ↘ |

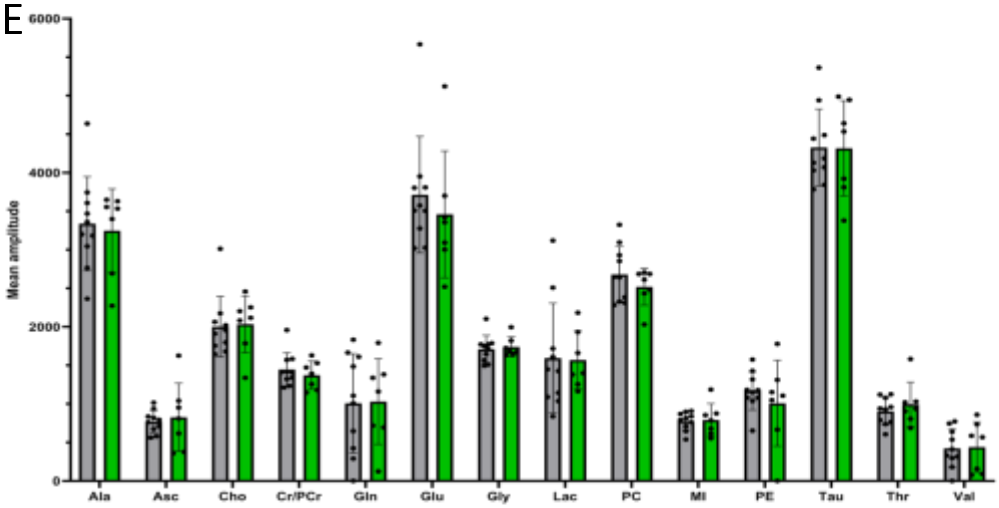


Fig. S4 Related to Fig. 4

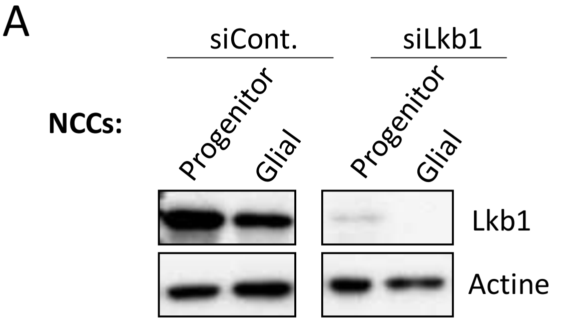


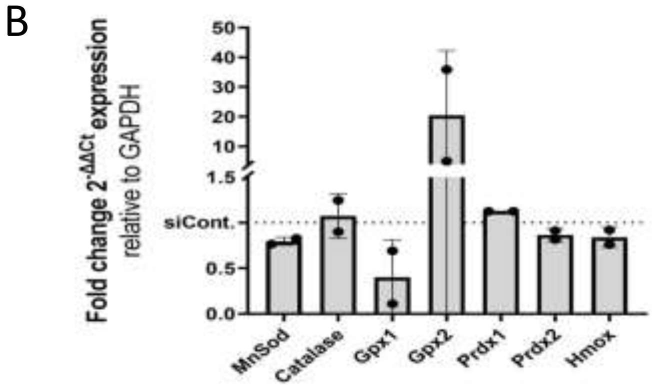


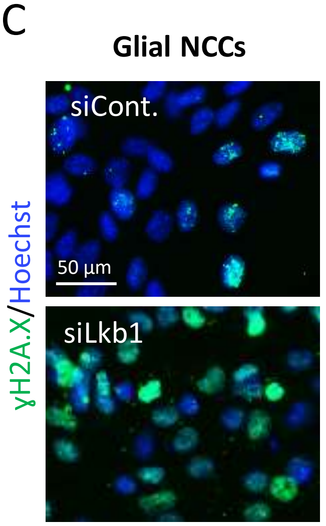


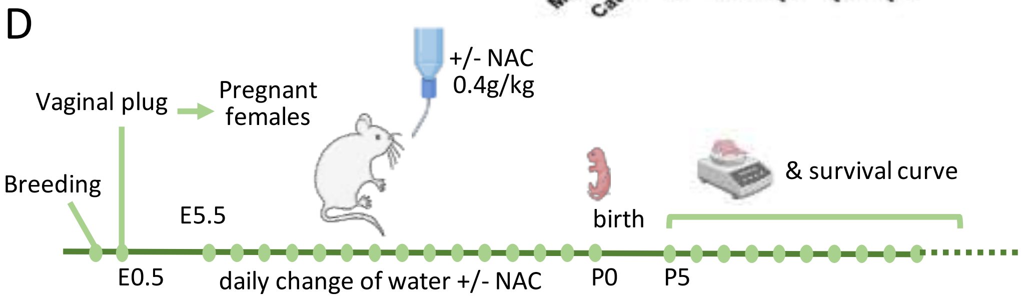


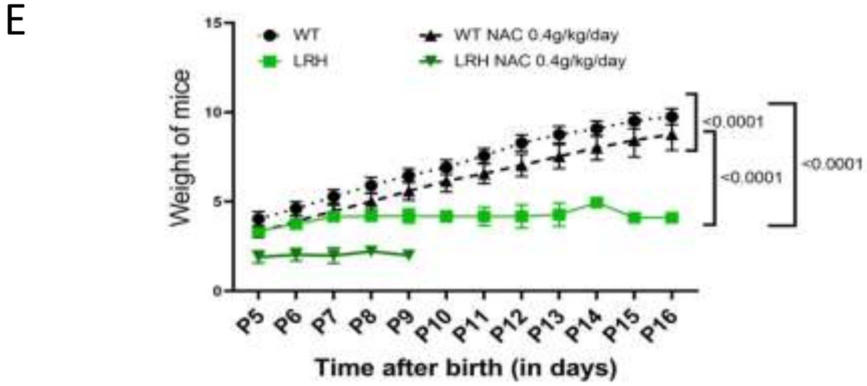


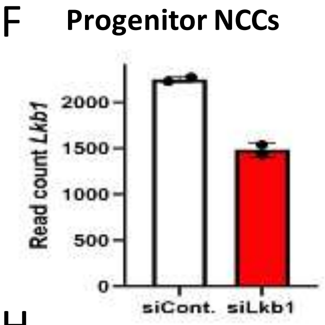


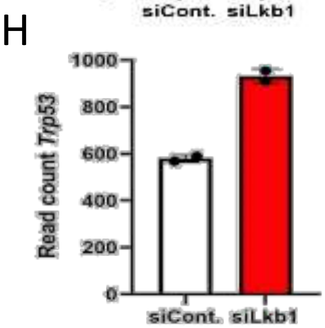


G

| GO term pathway | Total genes in the pathway | Rich ratio | Q value | Gene symbols |
|---|---|---|---|---|
| Apoptotic process (GO:0006915) | 627 | 0.008 | 0.036 | Stk11,Tia1,Trp53,Trib3, Chac1 |
| Circadian rhythm (GO:0007623) | 103 | 0.029 | 0.036 | Atf5,Trp53,Mat2a |
| Positive regulation of neuron projection development (GO:0010976) | 179 | 0.016 | 0.036 | Scarb2,Itga6,Gpc2 |
| Autophagy (GO:0006914) | 183 | 0.017 | 0.036 | Stk11,Trp53,Stbd1 |
| Glycogen metabolic process (GO:0005977) | 42 | 0.048 | 0.036 | Ppp1ccb,Stbd1 |
| Cell adhesion (GO:0007155) | 616 | 0.006 | 0.041 | Acan,Itga6,Msln,Rsc1a1 |
| Regulation of cell cycle (GO:0051726) | 279 | 0.011 | 0.041 | Atf5,Stk11,Trp53 |
| Central nervous system development (GO:0007417) | 117 | 0.017 | 0.041 | Acan,Trp53 |

Fig. S5. Related to Fig. 5

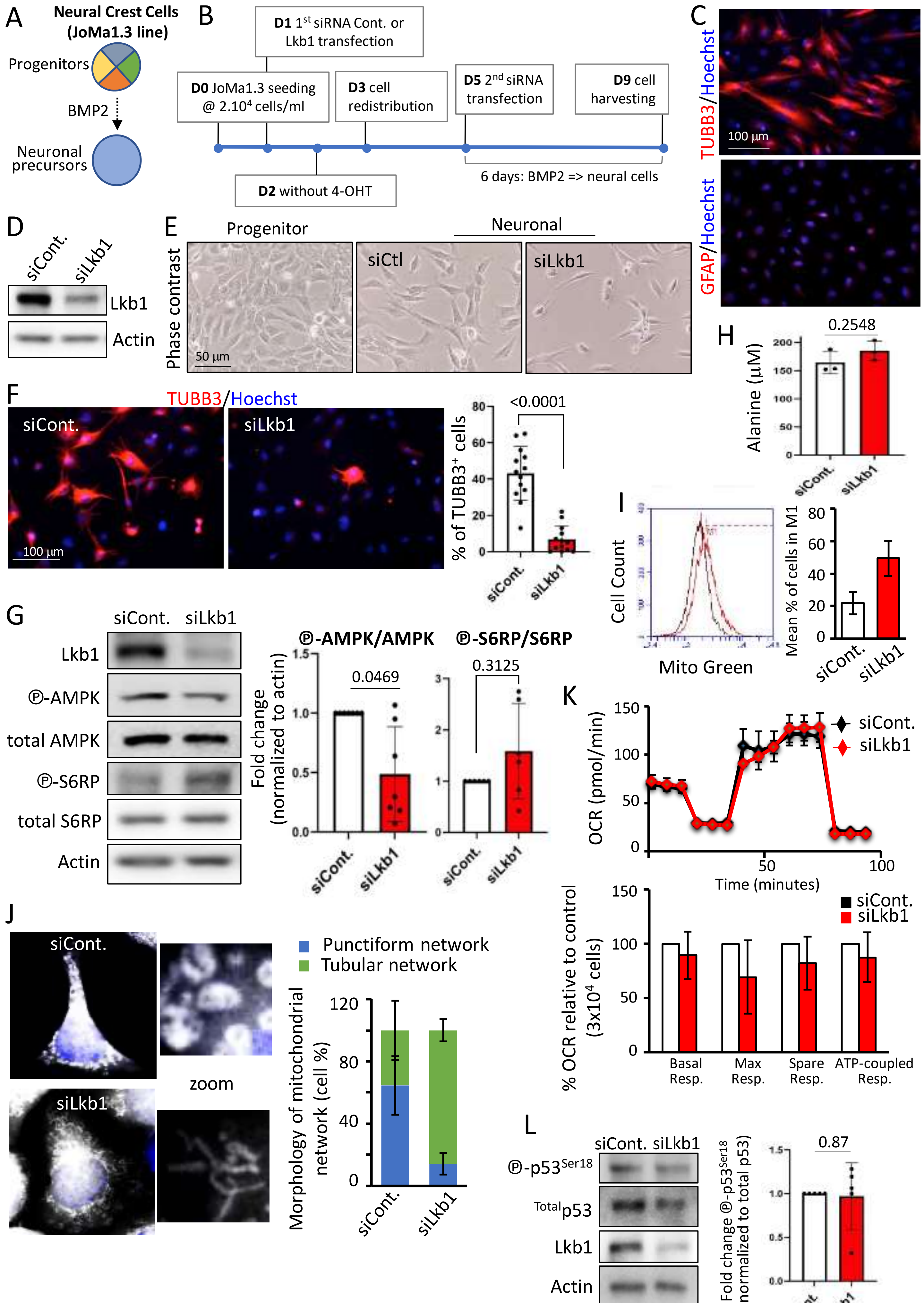


Fig. S6 Related to Fig. 5

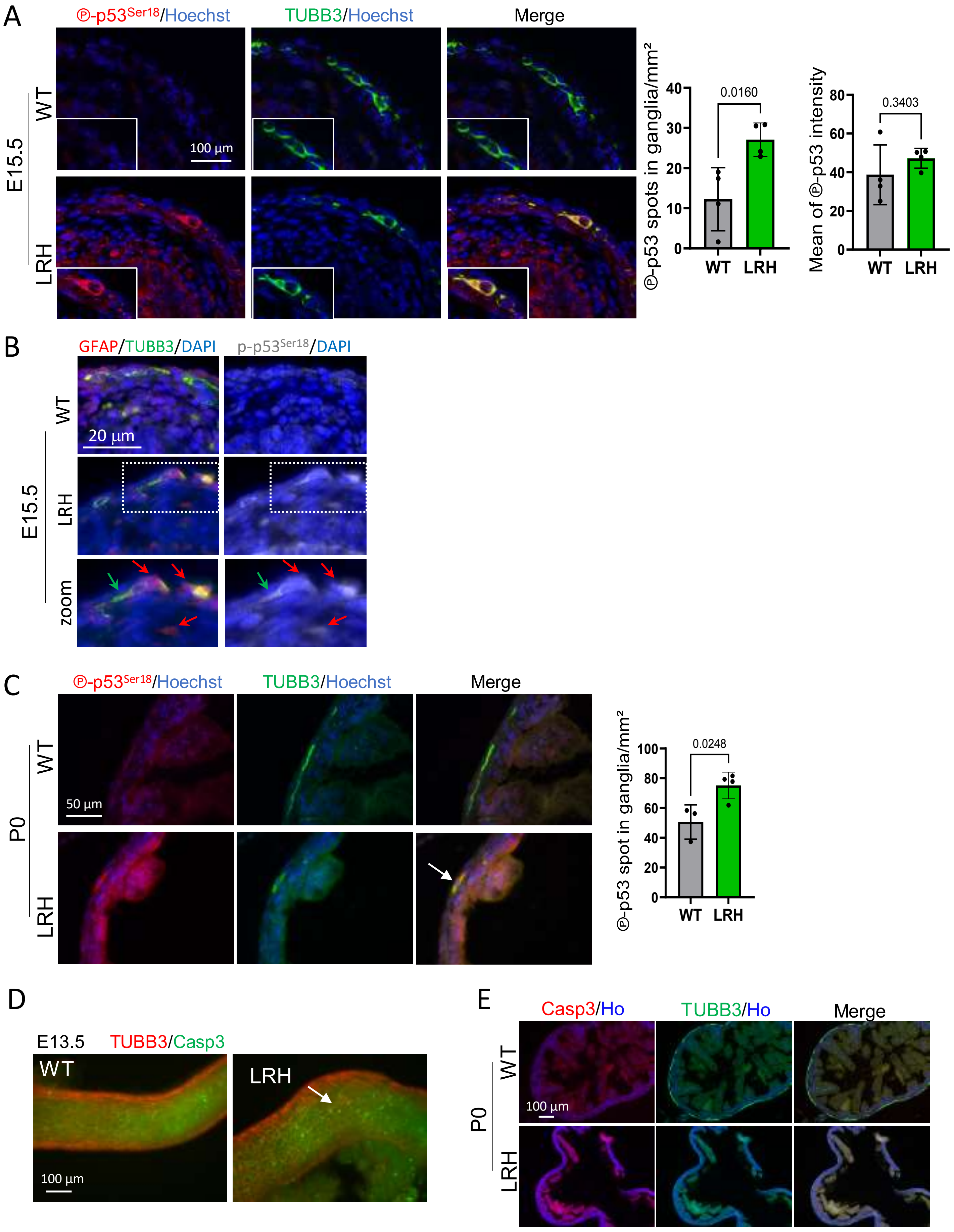


Fig. S7 related to Fig. 6

A

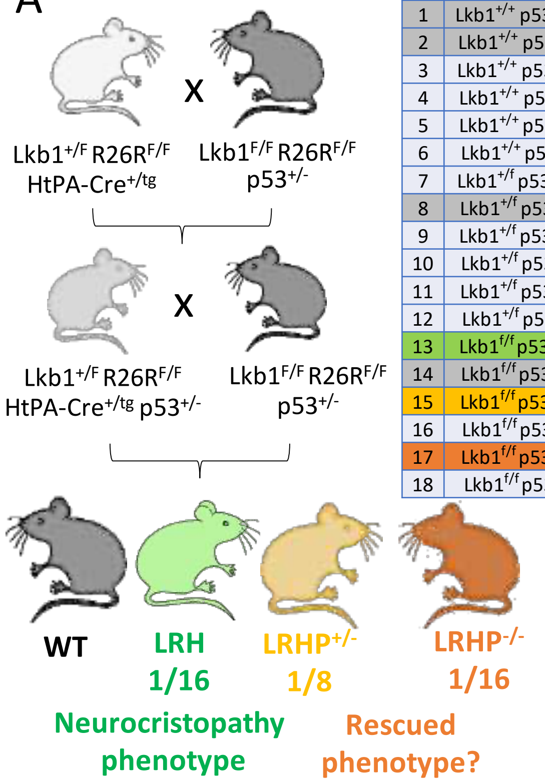


| N° | genotype |
|---|---|
| 1 | Lkb1$^{+/+}$ p53$^{+/+}$ HtPA-Cre$^{+/tg}$ |
| 2 | Lkb1$^{+/+}$ p53$^{+/+}$ HtPA-Cre$^{+/+}$ |
| 3 | Lkb1$^{+/+}$ p53$^{+/-}$ HtPA-Cre$^{+/tg}$ |
| 4 | Lkb1$^{+/+}$ p53$^{+/-}$ HtPA-Cre$^{+/+}$ |
| 5 | Lkb1$^{+/+}$ p53$^{-/-}$ HtPA-Cre$^{+/tg}$ |
| 6 | Lkb1$^{+/+}$ p53$^{-/-}$ HtPA-Cre$^{+/+}$ |
| 7 | Lkb1$^{+/f}$ p53$^{+/+}$ HtPA-Cre$^{+/tg}$ |
| 8 | Lkb1$^{+/f}$ p53$^{+/+}$ HtPA-Cre$^{+/+}$ |
| 9 | Lkb1$^{+/f}$ p53$^{+/-}$ HtPA-Cre$^{+/tg}$ |
| 10 | Lkb1$^{+/f}$ p53$^{+/-}$ HtPA-Cre$^{+/+}$ |
| 11 | Lkb1$^{+/f}$ p53$^{-/-}$ HtPA-Cre$^{+/tg}$ |
| 12 | Lkb1$^{+/f}$ p53$^{-/-}$ HtPA-Cre$^{+/+}$ |
| 13 | Lkb1$^{f/f}$ p53$^{+/+}$ HtPA-Cre$^{+/tg}$ |
| 14 | Lkb1$^{f/f}$ p53$^{+/+}$ HtPA-Cre$^{+/+}$ |
| 15 | Lkb1$^{f/f}$ p53$^{+/-}$ HtPA-Cre$^{+/tg}$ |
| 16 | Lkb1$^{f/f}$ p53$^{+/-}$ HtPA-Cre$^{+/+}$ |
| 17 | Lkb1$^{f/f}$ p53$^{-/-}$ HtPA-Cre$^{+/tg}$ |
| 18 | Lkb1$^{f/f}$ p53$^{-/-}$ HtPA-Cre$^{+/+}$ |

B

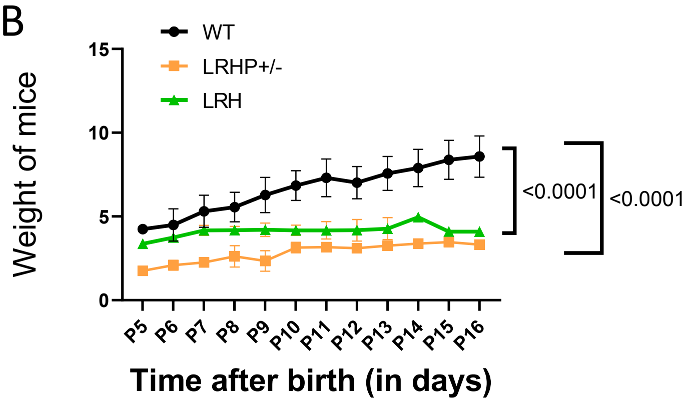


C

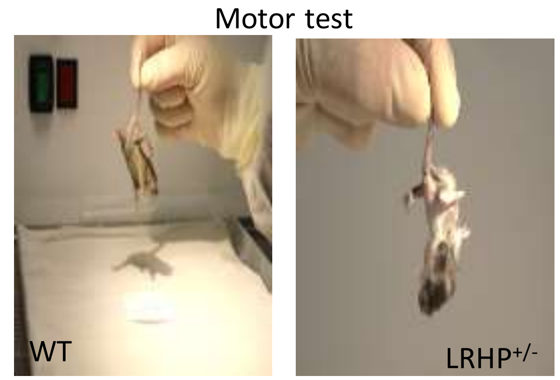


D

| Age | Nb of litters | LRH observed number (expected) | WT observed number (expected) | Chi² value |
|---|---|---|---|---|
| **E10.5** | 19 | 29 (26.5) | 24 (26.5) | **0.47 (ns)** |
| **E13.5** | 8 | 17 (18) | 19 (18) | **0.11 (ns)** |
| **E15.5** | 6 | 15 (11.5) | 8 (11.5) | **2.13 (ns)** |
| **All (10.5-15.5)** | **33** | **61 (56)** | **51 (56)** | **0.89 (ns)** |

E

| Questions asked | Involved genotypes | Nb of pups | Chi² p value | Answer |
|---|---|---|---|---|
| Global Mendelian inheritance ? | all, 1-18 | **703** | **$2.1\ 10^{-9}$** | **NO** |
| Mendelian inheritance in control genotypes ? | 7-10,14,16 | 534 | 0.09 | YES |
| Mendelian inheritance of p53$^{+/-}$ and p53$^{-/-}$ with Lkb1$^{+/+}$ ? | 1-6, 8,10,12,14,18 | 441 | 0.11 | YES |
| Mendelian inheritance of Lkb1 with p53$^{+/+}$ ? | 1,2,7,8,13,14 | **342** | **$5.3\ 10^{-7}$** | **NO** |
| No lethality of htz LRH mutants? | 7,8 | 199 | 0.98 | YES |
| No lethality of hmz LRH mutants? | 13,14 | **138** | **$7.9\ 10^{-7}$** | **NO** |
| Estimated lethality of Hmz LRH mutants | 13,14 | **138** | **58%** | |
| Mendelian inheritance if including 58% Hmz LRH lethality? | all, 1-18 | 703 | 0.26 | YES |

F

| Lkb1 & HtPA genotypes | Involved genotypes | Observed Nb. | Expected Nb. (Mendelian) | *Chi² and p value* | Estimated lethality | Theoretical Nb. (with lethality) | *Chi² and p value* |
|---|---|---|---|---|---|---|---|
| **WT** | 1,2,8,14 | 216 | 176.75 | *28.9* | -- | 204.21 | *2.29* |
| **Htz LRH** | 7 | 86 | 85.00 | *$p < 10^{-6}$* | -- | 98.55 | ***ns*** |
| **Hmz LRH** | 13 | 40 | 80.25 | | **58%** | 39.24 | |

Fig. S8 Related to Fig. 7

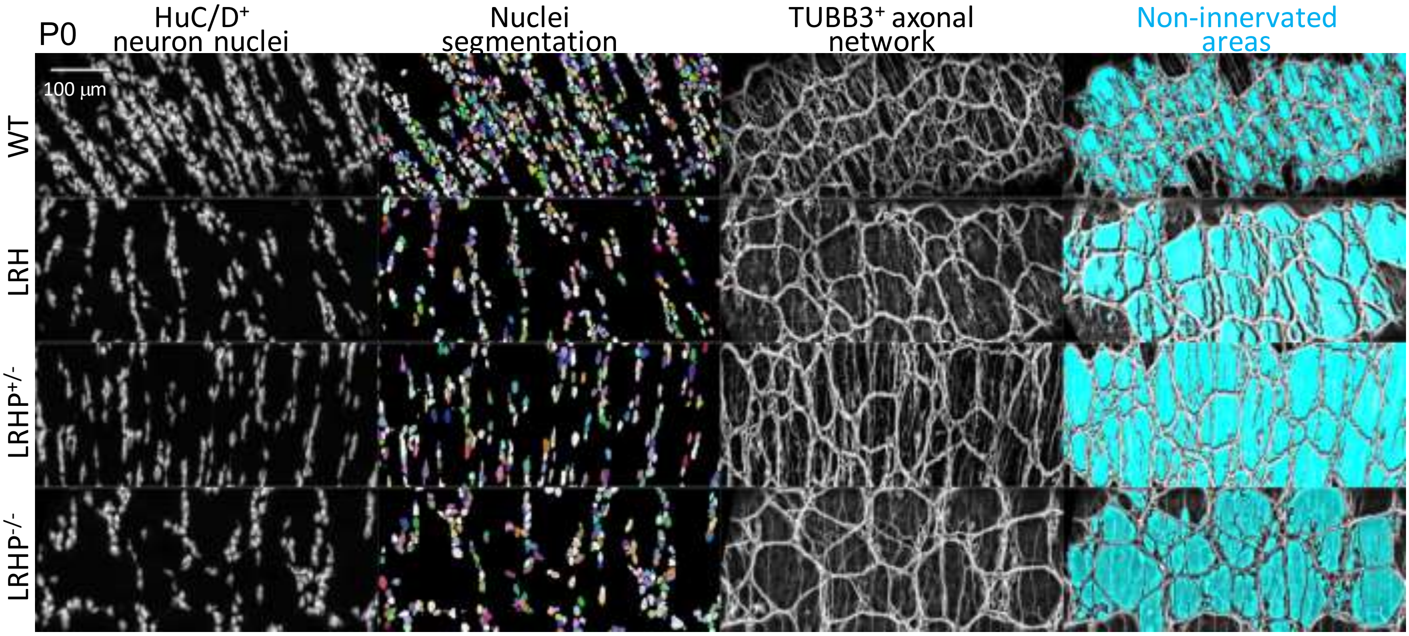


Fig. S9 Related to Fig. 7
